# Nucleoid-associated proteins sense phage-induced genome damage to elicit abortive infection

**DOI:** 10.64898/2026.04.06.716658

**Authors:** Zhihua Li, Shuai Yuan, Jiayue Ma, Xian Shu, Julian J. Duque-Pedraza, Ilya Terenin, Zonglan Yu, Feiyue Cheng, Jinhui Wang, Abel Garcia-Pino, Gemma Atkinson, Hang Yang, Jingmin Gu, Vasili Hauryliuk, Shouyue Zhang, Bing Liu, Ming Li

**Affiliations:** CAS Key Laboratory of Microbial Physiological and Metabolic Engineering, State Key Laboratory of Microbial Diversity and Innovative Utilization, Institute of Microbiology, Chinese Academy of Sciences, Beijing, China; College of Life Science, University of Chinese Academy of Sciences, Beijing, China; Key Laboratory of Virology and Biosafety, Wuhan Institute of Virology, Chinese Academy of Sciences, Wuhan, 430071, China; NanoLund, Lund University, Lund, Sweden; Department of Infectious Diseases, Key Laboratory of Surgical Critical Care and Life Support, BioBank, The First Affiliated Hospital of Xi’an Jiaotong University, Xi’an Jiaotong University, Shaanxi, 710061, China; Cellular and Molecular Microbiology, Faculté des Sciences, Université Libre de Bruxelles (ULB), Brussels, Belgium; Science for Life Laboratory, Lund, Sweden; State Key Laboratory for Diagnosis and Treatment of Severe Zoonotic Infectious Diseases, Key Laboratory for Zoonosis Research of the Ministry of Education, Institute of Zoonosis, and College of Veterinary Medicine, Jilin University, Changchun 130062, China; University of Tartu, Institute of Technology, Tartu, Estonia; Beijing Key Laboratory of Genetic Element Biosourcing & Intelligent Design for Biomanufacturing, Beijing 100101, China

## Abstract

Bacteria have evolved diverse immune strategies to detect and neutralize bacteriophage infection. Here, we describe an unprecedented paradigm in which a chromosome-architecting nucleoid-associated protein (NAP) is repurposed as a viral infection sensor. When phage attack leads to genome degradation, the NAP sensor is released from the nucleoid to the cytoplasm, where it binds and activates diverse immune effectors. One such effector is a nucleotide-modifying toxin normally existing as an inactive homotetramer. NAP binding converts it into a catalytically active heterotrimer that halts both transcription and translation. Phylogenetic analyses unveiled the high modularity, polyphyletic origin, and wide distribution of NAP-mediated defenses. Collectively, we define a distinct class of defense systems in which bacteria sense phage-induced genome damage through NAP relocation, highlighting an unexpected but essential role for these proteins as sentinels of genome integrity.

---

Bacteria and their viruses (bacteriophages) have engaged in an intense evolutionary arms race for billions of years. Phages pose a significant threat to bacterial survival by hijacking cellular machinery to replicate and propagate^1^. In response to this pressure, bacteria have evolved a diverse array of anti-phage defense systems^2^. These systems either provide cellular-level immunity by specifically targeting phages to rescue the infected cells or act on the population level by sacrificing infected cells to prevent phage proliferation, thereby saving the rest of the population^3–6^.

The first and critical step in all anti-phage defense mechanisms is the detection of infection. A common strategy is directly sensing of viral components such as DNA, RNA or proteins. Discriminative detection of phage DNA typically relies on host DNA modifications, such as methylation employed by restriction-modification (RM) systems and less-characterized RM-like systems (e.g., BREX and DISARM)^7,8^. If phage nucleic acids evade detection, phage proteins will be actively synthesized. Defense systems like Avs (antiviral systems), Stk2, DSR (defense-associated sirtuins), and fused toxSAS toxin-antitoxin CapRel directly sense phage proteins essential for replication or assembly, such as terminases, portal proteins or capsid proteins^9–12^. Upon detection, these protein-sensing systems typically activate toxic effectors, leading to cell death and blocking phage propagation.

In addition to direct detection of viral components, phage infection can be sensed by monitoring the integrity of host processes or components that are hijacked or destroyed by phages^13^. For example, retron systems like Ec48 detect phage-induced inhibition of the RecBCD complex^14^. Diverse toxin-antitoxin (TA) systems, such as ToxIN and RnlAB, monitor changes in cellular transcriptional activity: when phage-induced inhibition of host transcription precludes the synthesis of labile antitoxins, active toxins are released, thereby triggering altruistic cell death to protect the population^15,16^. While defenses that directly detect viral components are relatively well studied, the mechanisms that monitor bacterial cellular machinery remain poorly understood.

In the early stages of infection, phages commonly deploy nucleases to degrade the host bacterial genome, thereby promoting viral DNA replication and enhancing progeny virion production^17,18^. Here, we describe PANGU – named after a Chinese deity – a distinct group of highly modular and widely distributed bicistronic immune systems encoding a nucleoid-associated protein (NAP) that clamps the bacterial chromosome. Focusing on PANGU variants encoding an alarmone-synthesizing toxSAS effector Pag1B/GRel, we show that the NAP sensor is released from the nucleoid when phage nucleases degrade bacterial genome, migrates to the cytoplasm and triggers the toxin effector. In the inactive state, the toxin forms an inert homotetramer. Upon NAP binding, this tetramer switches stoichiometry to an active heterotrimer, which actively catalyze the conversion of ATP and GTP to AMP and the alarmone guanosine pentaphosphate (pppGpp), thereby halting essential cellular pathways to abort the infection. Our bioinformatic analyses uncovered at least 10 distinct types of putative PANGU-like defense systems, which distribute across a variety of bacterial phyla and exhibit high modularity and a polyphyletic origin. This study establishes a distinct paradigm for phage infection detection: immunity is triggered not by binding a specific viral molecule, but by sensing virus-induced genome degradation and converting this cue into the relocation of a chromosome-bound sensor that, once displaced, licenses toxin activation. Collectively, our work reveals – unprecedentedly – the coordinated sensing and aborting phage infection by chromosome-architecting NAPs and nucleotide-modifying toxins.

## Two types of PANGU defend *E. coli* against coliphages

Despite YejK being a highly abundant NAP in *Escherichia coli*, its physiological role still remains poorly understood^19,20^. While conducting our manual search for YejK-like NAP proteins across bacterial genomes, we identified a number of NAP-encoding operons with genetic linkage to known anti-phage defense systems such as restriction-modification, Hachiman, Gabija and others (Extended Data Fig. 1). To test whether NAP-encoding operons also mediate defense, we synthesized a dozen codon-optimized operons and assessed their antiviral activity in *E. coli*. Four of the tested systems show pronounced anti-phage effects against T4 or Bas34, reducing their viral plaque-forming units (PFUs) by 3-4 logs (Fig. 1a, b). We defined such NAP-related bicistronic (two-gene) anti-phage operons as PANGU. Type I PANGU operons encode a NAP protein (designated Pag1A) and small alarmone synthetase (toxSAS) toxin Pag1B that belongs to unexplored GRel toxSAS sub-family^21,22^ (Fig. 1a). toxSAS toxins target either translation via tRNA pyrophosphorylation or cellular metabolism via synthesis of alarmone nucleotides^12,21,22^. Type II PANGU encode a NAP protein Pag2A together with a transmembrane (TM) protein effector Pag2B (Fig. 1b).

**Fig. 1.**
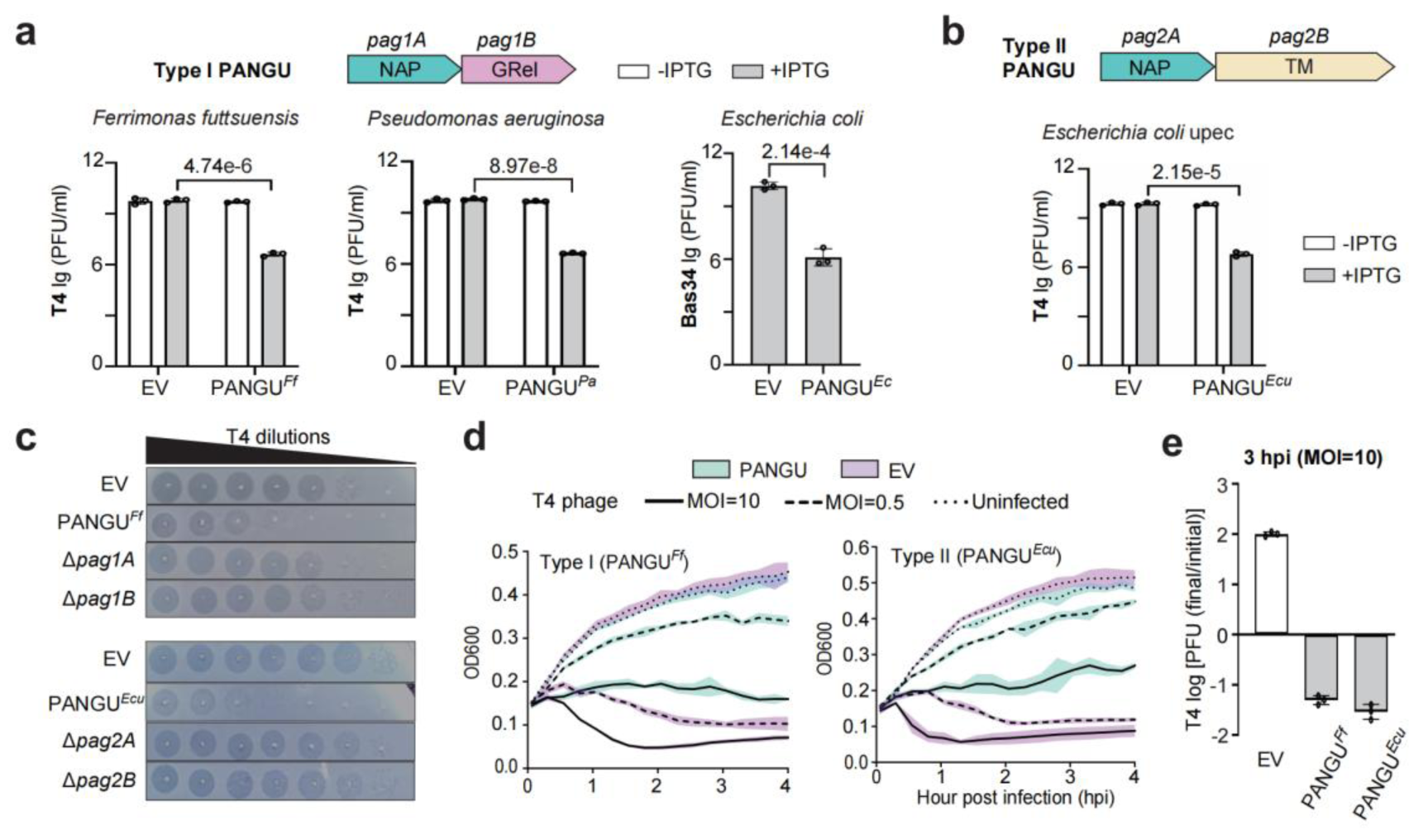
Two types of PANGU systems defend *E. coli* against phages. **a**, **b**, Infectivity of T4 or Bas34 infecting *E. coli* cells expressing a type-I (**a**) or type-II (**b**) PANGU variant expressed in Plaque forming units (PFUs). EV, empty vector. PANGU*^Ec^* was under the control of a constitutive promoter, while IPTG was used to induce the other three operons (see Methods). *P* values were calculated from two-sided Student’s *t* test. **c**, Serial tenfold dilutions of T4 spotted on lawns of *E. coli* cells expressing either the intact bicistronic PANGU*^Ff^* or PANGU*^Ecu^* systems, or the individual *pagA* or *pagB* genes. **d**, Growth curves of PANGU-expressing *E. coli* cells infected with T4 at increasing multiplicities of infection (MOIs). **e,** Proliferation of T4 phage during a 3-hour infection at an initial MOI of 10. *E. coli* cells expressing PANGU*^Ff^* or PANGU*^Ecu^* were infected and cultures were harvested at 3 hours post-infection. PFUs per milliliter of supernatant were quantified on control cell lawns, and total PFUs were calculated and divided by the initial PFUs used for infection. Data represent the mean ± SD of three biological replicates.

To probe the importance of PagA and PagB in defense, we subjected the Type I PANGU system from *F. futtsuensis* (PANGU*^Ff^*) and the Type II operon from *Escherichia coli* upec (PANGU*^Ecu^*) for gene deletion analysis. For PANGU*^Ff^*, both Pag1A and Pag1B are essential for phage resistance (Fig. 1c). Similarly, both Pag2A and Pag2B were critical in the case of PANGU*^Ecu^* (Fig. 1c). Importantly, none of the constructs were toxic, suggesting that unlike the toxSAS-encoding systems studied before, PANGU is not a classical toxin-antitoxin system^21^; rather, PagB might act as a phage sensor that licenses the PagA activity once triggered by the virus.

To determine whether PANGU systems confer protection at the individual or population level, we performed T4 infection experiments at low (0.5) or high (10) multiplicity of infection (MOI) (Fig. 1d). At MOI 0.5, while the culture expressing either PANGU*^Ff^* or PANGU*^Ecu^* systems did not collapse and could grow despite the infection, we did not observe complete immunity; the growth was stunted as compared to uninfected controls. At MOI 10, while both systems prevented phage-induced culture collapse, bacterial growth was impaired even more severely, with cultures exhibiting either stagnation or extremely slow proliferation (Fig. 1d). The absence of complete immunity at low MOIs and the extremely slow growth at high MOIs suggest that PANGU acts through abortive infection, whereby infected cells become non-replicative due to phage-triggered toxicity. Consistently, the titer of extracellular infectious phages in PANGU-expressing cultures decreased by 1.5 logs within 3 hours post-infection, in contrast to the 2-log increase observed in empty vector control (Fig. 1e). Collectively, these results indicate that PANGU restricts phage propagation at the population level by limiting the viability of infected cells and thereby reducing viral spread.

## Pag1A-Pag1B complex produces the alarmone pppGpp

We next explored how Type I PANGU limits the viability of infected cells. AlphaFold^23^ modeling of Pag1B/GRel toxin ^21^ encoded by PANGU*^Ff^* revealed that the protein is made up an N-terminal enzymatic toxSYNTH domain and a C-terminal domain of unknown function bridged by an α-helical linker (Fig. 2a and Extended Data Fig. 2a). The C-terminal domain of Pag1B has an acidic, surface-enriched composition mimicking the properties of DNA; we have termed it a DNA-mimicking (DMK) domain. The best-studied antiphage toxSAS is CapRel^SJ^^46^, a fused translation-targeting tRNA-modifying toxin-antitoxin from *Salmonella* phage SJ46^24^. Structural comparison between CapRel^SJ^^46^ and the N-terminal toxSYNTH domain of Pag1B reveal that the two share a common catalytic fold (RMSD = 2.73 Å; Fig. 2b). However, unlike CapRel^SJ^^46^, Pag1B lacks the positively charged pocket that is inveigled in tRNA binding, indicating its potential function as a metabolism-targeting toxin involved in alarmone production. (Extended Data Fig. 2b). The conserved G-loop tyrosine (Y155 in CapRel ^SJ^^46^) required for enzymatic functionality^12^ corresponds to Y129 in Pag1B. Replacing this tyrosine with alanine (Y129A) abolished the PANGU*^Ff^*-mediated defense against T4 phage (Fig. 2c), confirming that the catalytic activity of Pag1B is indispensable for immunity.

**Fig. 2.**
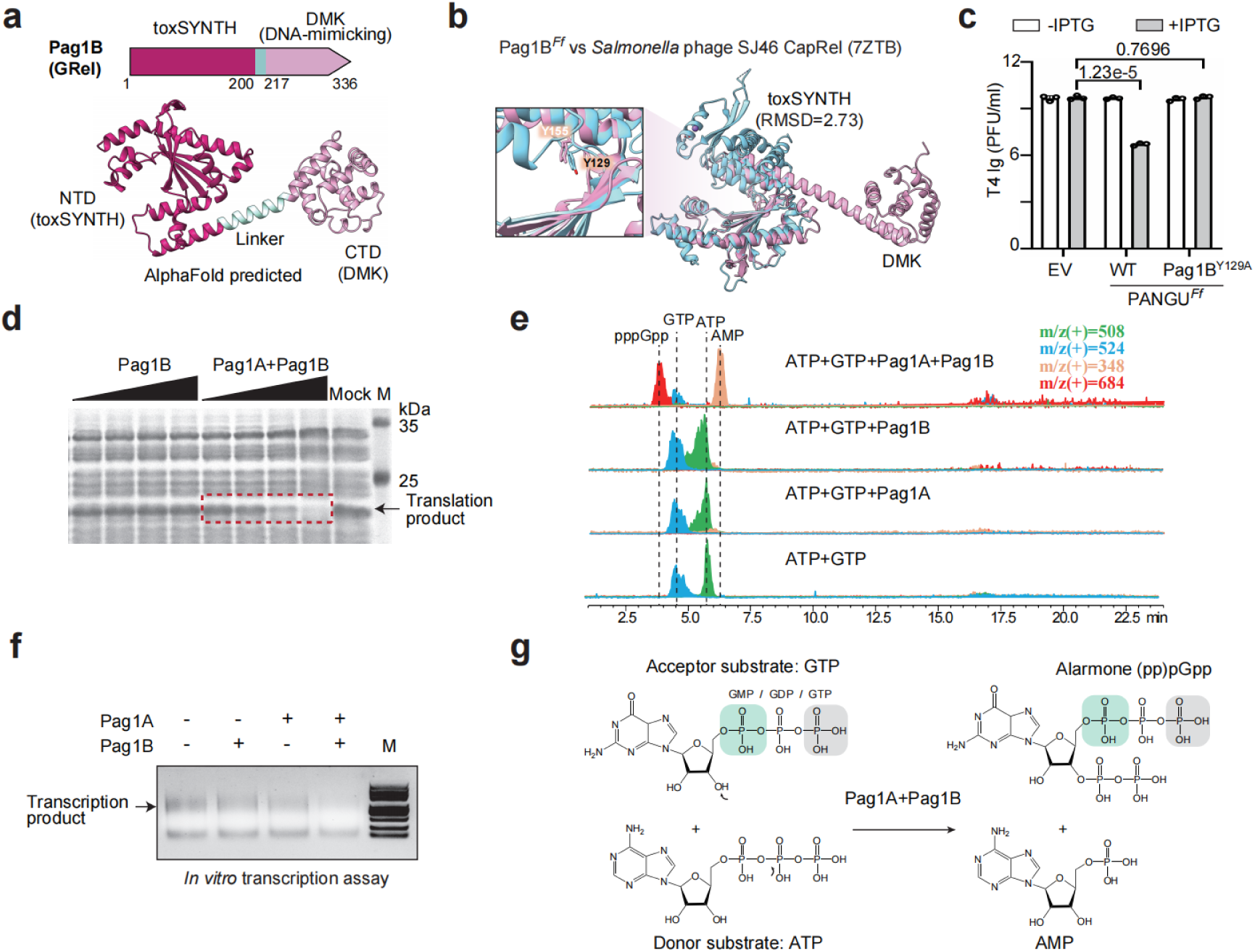
Pag1A-activated Pag1B converts ATP and GTP to AMP and pppGpp. **a,** AlphaFold-predicted structure of the GRel toxSAS Pag1B (see Extended Data Fig. 2a). **b,** Superposition of Pag1B (pink) onto CapRel^SJ^^46^ (light blue, PDB 7ZTB). The root-mean-square deviation (RMSD) value for the alignment of their toxSYNTH domains is indicated. The boxed enlargement shows the superposition of catalytic tyrosines (Y155 in CapRel ^SJ^^46^ and Y129 in Pag1B). **c,** PFUs of T4 infecting *E. coli* cells expressing either wild-type PANGU*^Ff^* or a PANGU*^Ff^* variant with catalytically-dead Pag1B^Y129A^. IPTG was used for induction. Data represent the mean ± standard deviation (SD) of three biological replicates. **d,** Inhibitory activity of PANGU*^Ff^* assessed in a coupled *in vitro* transcription-translation assay using DHFR production from a DNA template as the readout. T7 RNA polymerase was used. Purified Pag1B was tested alone or with Pag1A, each at concentrations of 62.5, 125, 250, and 500 nM. Mock, protein storage buffer. M, protein markers in kilodaltons (kDa). **e,** LC-MS profiles of the products formed from ATP and GTP after incubation with purified Pag1A and/or Pag1B. **f**, *In vitro* transcription assay using purified *E. coli* RNA polymerase. M, 100-bp DNA marker. Representative images from three independent biological replicates are shown. **g,** Scheme of the pyrophosphorylation reaction catalyzed by Pag1A and Pag1B.

To probe the toxic activity of Pag1B/GRel and the possible role of Pag1A in its activation, we first used an *in vitro* coupled transcription-translation system ^12^. When added alone, purified Pag1B did not suppress the synthesis of a control dihydrofolate reductase (DHFR) reporter (Fig. 2d). As expected, addition of Pag1A alone neither had inhibitory effects (Extended Data Fig. 3a). However, when Pag1A and Pag1B were added at an equimolar concentration, DHFR production was repressed, and signal disappeared completely at 500 nM each of Pag1A and Pag1B (Fig. 2d). We next asked whether inhibition is achieved through tRNA pyrophosphorylation. Incubation of Pag1A/Pag1B with *E. coli* total tRNA and [γ³²P]ATP revealed no radiolabel transfer (Extended Data Fig. 3b), ruling out this mechanism and suggesting that the inhibitory effect is indirect and likely mediated by the alarmone production^25,26^.

To probe the alarmone-synthesizing activity, when ATP and GTP were supplied to Pag1A-activated Pag1B, the nucleotides were converted to AMP and pppGpp and the NTP substrates were depleted (Fig. 2e). None of the other tested other NTPs or dNTPs could serve as substrates (Extended Data Fig. 3c, d). Importantly, Pag1A-activated Pag1B can also pyrophosphorylate GMP and GDP, producing ppGpp and pGpp alarmones (Extended Data Fig. 3e). Consistent with NTP depletion and the accompanying rise in alarmone levels, transcription by purified *E. coli* RNA polymerase was markedly inhibited by Pag1A and Pag1B (Fig. 2f). Therefore, unlike CapRel, which targets tRNAs, Pag1A-stimulated GRel toxin Pag1B uses ATP-derived pyrophosphate to convert GTP/GDP/GMP into (pp)pGpp (Fig. 2g), thereby globally suppressing transcription, translation, and likely other essential processes, to abort infection.

## Pag1A and Pag1B form a heterotrimer

We next asked how Pag1A activates Pag1B. Given that the C-terminal domain of Pag1B exhibits an acidic, surface-enriched composition that mimics DNA properties (Fig. 3a), we first assessed whether the two proteins engage in a direct physical interaction. A pull-down assay showed that Pag1B, indeed, could be captured by Flag-tagged Pag1A (Fig. 3b). Furthermore, bio-layer interferometry (BLI) assay showed that the interaction is exceedingly tight, as Pag1A binds to DMK with a high affinity (*KD* = 9.4 nM) (Fig. 3c).

**Fig. 3.**
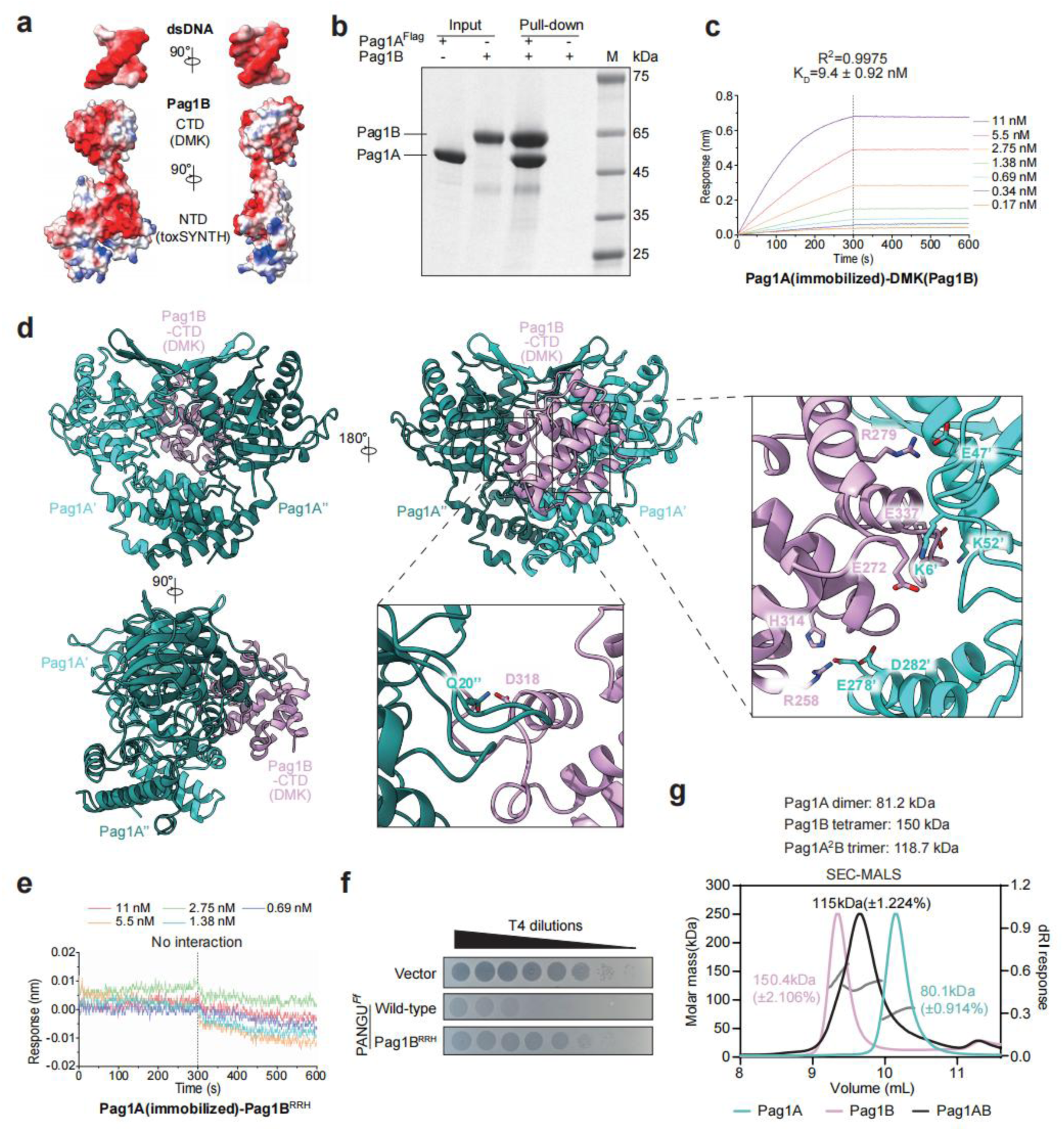
Pag1A and Pag1B form a heterotrimeric complex. **a,** Comparing the surface charge distribution of Pag1B and a random double-stranded DNA (dsDNA) fragment using electrostatic potential mapping. Surface atoms are color-coded in a red-to-blue gradient, ranging from negative to positive charges, respectively. The visualization was generated using ChimeraX and displayed with a 90° rotational offset along the z-axis to highlight structural and electrostatic similarities between the two molecules. **b,** *In vitro* pull-down of Pag1B using Flag-labeled Pag1A as a bait. M, protein markers in kilodaltons (kDa). **c,** Bio-Layer Interferometry (BLI) analysis of Pag1A and the C-terminal DMK domain of Pag1B. Serial dilutions of the DMK domain of Pag1B were injected onto Pag1A-loaded biosensors for a 300-second association phase, followed by a 300-second dissociation phase. **d,** The cryo-EM structure of the Pag1A-Pag1B complex was determined at 2.64 Å resolution, which reveals a heterotrimeric assembly composed of two Pag1A subunits (Pag1A′ and Pag1A″) and the C-terminal DMK domain of Pag1B. The N-terminal toxSYNTH domain of Pag1B could not be detected likely due to flexibility in the linker region. **e,** BLI analysis of Pag1A and the Pag1B^RRH^ mutant (residues R258, R306, and H314 that interact with Pag1A were simultaneously substituted with an alanine). **f,** T4 infectivity on *E. coli* cells expressing wild-type PANGU*^Ff^* or its mutant with Pag1B^RRH^. Representative images from three independent biological replicates are shown. **g,** SEC-MALS analysis of purified *P. aeruginosa* Pag1A, Pag1B, or an equimolar mixture (separately purified and subsequently co-incubated). The molecular weights of Pag1A and Pag1B monomers are indicated on the top.

Next, to elucidate the structural basis underlying the Pag1A-Pag1B interaction, we employed single-particle cryo-electron microscopy (cryo-EM), resolving the complex at an overall resolution of 2.64 Å (Extended Data Fig. 4). Pag1A and Pag1B form a heterotrimeric complex comprising two subunits of Pag1A (designated Pag1A ′ and Pag1A″) and the C-terminal DMK domain of Pag1B (Fig. 3d). Notably, while a full-length Pag1B protein was used for complex preparation, the N-terminal domain (NTD) of Pag1B was not detected in the final structure. This likely stems from the inherent flexibility of the linker between the NTD and CTD domains of Pag1B, or from interruption of the linker during sample preparation. (Fig. 2a).

The DMK domain of Pag1B mediates asymmetric interactions with the two Pag1A subunits. While it forms extensive contacts with Pag1A′, the interaction with Pag1A″ is relatively weak (Fig. 3d). The primary Pag1A′-Pag1B interface spans approximately 1,200 Å² and comprises six salt bridges connecting residues E47, K6, K52, D282, E278, and E278 of Pag1A to R279, E272, E337, R258, R258, and H314 of Pag1B, respectively, along with 12 hydrogen bonds and 144 non-bonded contacts. Conversely, the secondary Pag1A″-Pag1B interface is much smaller (only 400 Å²) and includes just one hydrogen bond and 17 non-bonded contacts (Fig. 3d). This pronounced asymmetry seemingly arises from the intrinsic structural asymmetry of Pag1B itself: while two copies of Pag1A are required to capture one Pag1B molecule, one Pag1A subunit is functionally specialized to assist the other in forming robust interactions with the DMK domain of Pag1B.

To disrupt one of the Pag1A′-Pag1B interfaces, we introduced three substitutions (R258A, R306A and H314A) into the DMK domain of Pag1B. BLI assays with the resultant triple-substituted Pag1B^RRH^ variant showed a complete loss of Pag1A binding (Fig. 3e). Furthermore, Pag1A and Pag1B^RRH^ pair neither inhibited DHFR synthesis (Extended Data Fig. 3a) nor conferred protection against T4 (Fig. 3f). Collectively, these results show that complex formation between Pag1A and Pag1B is indispensable for induction of Pag1B enzymatic activity and for the antiviral function of PANGU*^Ff^*.

Finally, we also analyzed the Type II PANGU*^Ecu^* complex. The AlphaFold-predicted structure of Pag2B revealed a C-terminal TM domain and an N-terminal DMK domain mimicking a longer DNA double helix, with a high density of acidic residues on its surface (Extended Data Fig. 2c). BLI analysis confirmed a high-affinity interaction between the isolated DMK domain and Pag2A (*KD* = 19 nM) (Extended Data Fig. 2d), suggesting that recognition between the core NAP protein and the cognate DMK domain is an architectural principle conserved across PANGU types.

## Pag1B forms an inactive tetramer

All toxSAS toxins studied to date function as toxic components of toxin-antitoxin systems, and their activity is inhibited by antitoxins acting either *in trans* (provided as separate polypeptides) or *in cis* (encoded within the same polypeptide)^15,16,21^. The Pag1A-Pag1B pair challenges this paradigm, as Pag1A functions as an activator rather than a classical antitoxin^27^. The fused toxin-antitoxin architecture of CapRel^12,24^ and the nanomolar affinity between Pag1A and the C-terminal DMK module of Pag1B (Fig. 3c) raised the possibility that the DMK module might act as an internal antitoxin. We therefore expressed the isolated enzymatic toxSYNTH domain of Pag1B. However, the fragment lacks the inhibitory activity (Extended Data Fig. 3a), ruling out this possibility and pointing to a different mechanism of triggering.

As Pag1B tended to aggregate during size-exclusion chromatography (SEC) (Extended Data Fig. 5a), we switched to a homologous Type I PANGU system from *Pseudomonas aeruginosa* (PANGU*^Pa^*) which similarly blocks phage proliferation (Fig. 1a) and inhibits transcription-translation system *in vitro* (Extended Data Fig. 5b). SEC revealed that purified *P. aeruginosa* Pag1B eluted as a discrete oligomer, which partially disassembled after co-incubation with its cognate Pag1A (Extended Data Fig. 5c, d). To precisely resolve the stoichiometry, we subjected the samples to SEC coupled with multi-angle light scattering (SEC-MALS). Pag1A (40.6-kDa) is a stable dimer (∼80.1 kDa), whereas the Pag1B alone (37.5 kDa) assembles into a tetramer (∼150.4 kDa) (Fig. 3g). Addition of the separately purified Pag1A disassembled the Pag1B tetramer and yielded a single 115-kDa species (Fig. 3g), precisely matching the molecular weight of the 2:1 heterotrimer observed in our crystal structure (Fig. 3d). Native PAGE revealed that the Pag1B tetramer progressively dissociated upon increasing addition of Pag1A (Extended Data Fig. 5e). Therefore, as revealed by the AlphaFold model (Extended Data Fig. 2a and Extended Data Fig. 5f), the ATP-binding pocket required for pyrophosphate transfer in the Pag1B tetramer appears to be sequestered at the subunit interface, and its binding to Pag1A may lead to the exposure of this pocket. Collectively, these data support a model in which inactive Pag1B forms a tetramer, and binding of Pag1A disassembles the tetramer to yielding the activated heterotrimeric immune effector.

## T4 escapes PANGU immunity though mutations in EndoII

Although the Pag1A-Pag1B complex is enzymatically active *in vitro* (Fig. 2e), co-expression of the two proteins in *E. coli* does not impair the growth: in the absence of phage infection, PANGU*^Ff^*-expressing cells grow similarly to the empty-vector control strain (Fig. 1d). Using two inducible promoters, we confirmed that expression of either or both proteins did not cause a growth defect on agar plates or in liquid medium (Extended Data Fig. 6a, b). This suggests that the catalytically-active Pag1A-Pag1B heterotrimer can only be formed upon phage infection.

To identify the trigger, we isolated T4 mutants that escape PANGU*^Ff^*-mediated defense. Using an experimental evolution approach^28^ (Extended Data Fig. 6c), we obtained four escape mutants (EM1-4) that produced PFUs on PANGU*^Ff^*-expressing cells comparable to those on vector-control cells (Fig. 4a). Genome sequencing revealed that each mutant carried a mutation in *denA*. This gene encodes endonuclease II (EndoII), which cleaves cytosine-containing host DNA but spares hydroxymethylcytosine-modified DNA of the T4 phage ^18^. EndoII functions as a constitutive tetramer with four catalytic centers^29^ (Fig. 4b). While EM1 carries a G29R substitution at the catalytic center, EM2 has C97W at the subunit-polymerization interface, with both potentially resulting in the loss-of-function (Fig. 4a, b). EM3 and EM4 harbor 28-bp and 1-bp frame-shift-causing deletions that introduce premature stop codons. As EM1-3 contain additional mutations elsewhere, we have primarily focused on EM4.

**Fig. 4.**
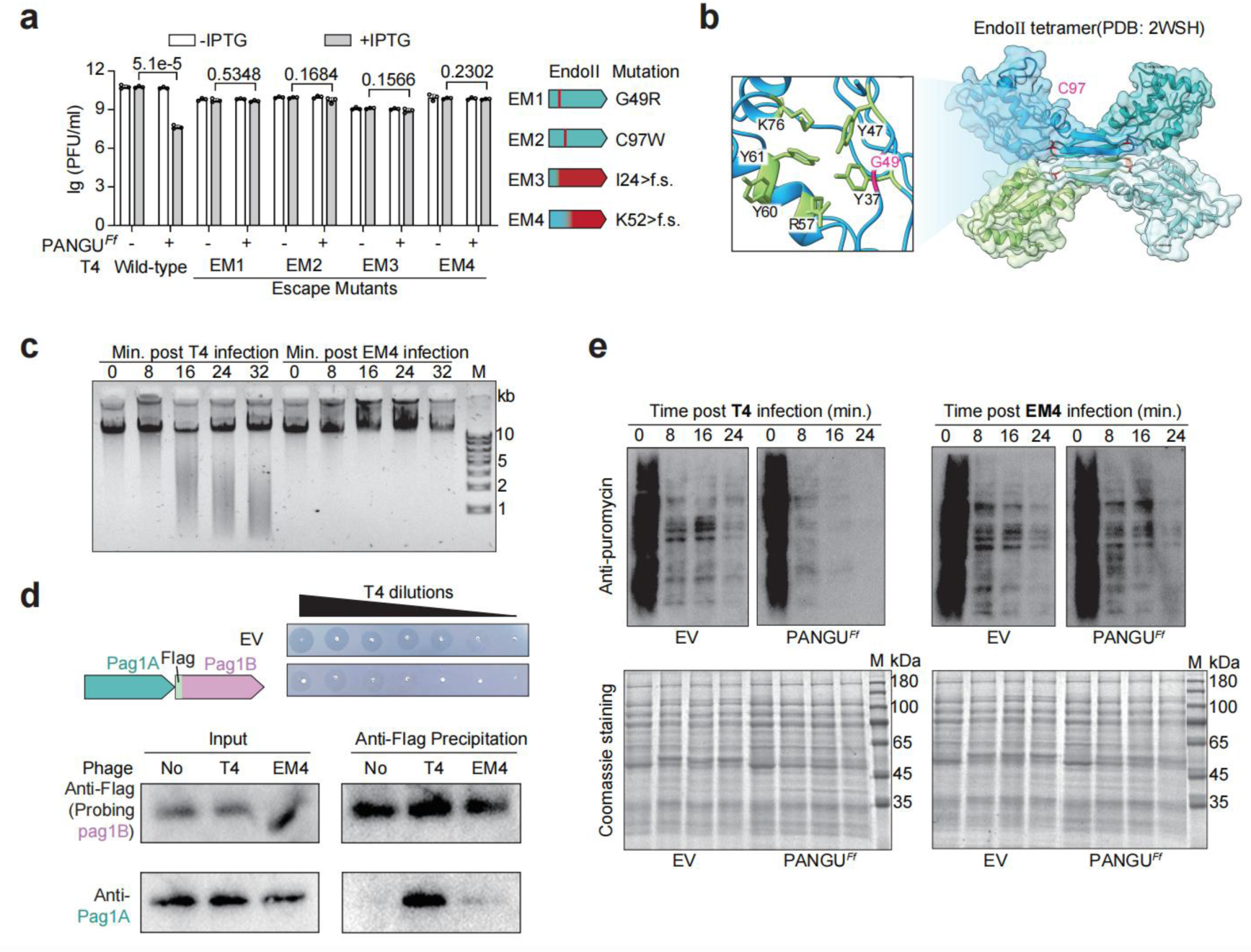
T4 EndoII is critical for induction of the Pag1A-Pag1B complex formation and activity. **a,** PFUs of wild T4 or its escape mutants (designated as EM1-4) infecting *E. coli* cells expressing PANGU*^Ff^* (the empty vector served as the negative control). IPTG was used for induction. Data represent the mean ± standard deviation (SD) of three biological replicates. Right show the different mutations in the *denA* gene (encoding EndoII) from EM1-4. f.s. : frame shift. **b,** The tetrameric structure of T4 EndoII highlighting the active residue G49 and the subunit-interacting residue C97. **c,** Agarose electrophoresis analysis of genomic DNA extracted from PANGU*^Ff^*-expressing *E. coli* cells infected by T4 or its EM4 mutant. M, 1-kb DNA marker. **d,** Co-immunoprecipitation of Pag1A using Flag-tagged Pag1B in *E. coli* cells under uninfected, wt-T4-infected, or EM4-T4-infected conditions. The immune function of this tagged PANGU*^Ff^* system was tested by spotting serial tenfold dilutions of T4 phages on the cell lawns. Pag1A was detected using anti-Pag1A antibodies. Representative images from three independent biological replicates are shown. **e,** Translational efficiency of PANGU*^Ff^-*expressing *E. coli* cells during infection with either wild-type T4 or EM4 T4 as monitored by puromycin incorporation and anti-puromycin immunoblotting. Representative images from three independent biological replicates are shown. Coomassie-stained total protein serves as loading control. M, protein markers in kilodaltons (kDa).

At the early stage of T4 infection, EndoII initiates host genome degradation, a process that also involves several viral endo- and exonucleases and ultimately generates nucleotide precursors for viral DNA replication^17,29^. Monitoring of genomic DNA integrity showed while infection with WT T4 results in extensive degradation already at 16 min post-infection, EM4 leaves the chromosome intact for at least 32 min, spanning a full T4 lytic cycle (Fig. 4c). Mutants EM1**–**3 similarly failed to degrade the *E. coli* genome (Extended Data Fig. 6d). Notably, expression of EndoII alone from a plasmid impaired *E. coli* growth (Extended Data Fig. 6e) and resulted in cleavage of the chromosomal DNA into ∼12-kb fragments but did not catalyze the extensive breakdown observed during the infection (Extended Data Fig. 6f). Our results are in line with the notion that while EndoII initiates host genome degradation, extensive degradation requires the subsequent action of other viral endo- and exonucleases^17,30,31^. However, no escape mutations were recovered in genes 46/47 or other ancillary nucleases, most likely because their products are essential for normal viral DNA replication^17,30,31^. Although EndoII is not essential, the *denA*-null EM4 mutant itself incurs a measurable fitness cost and is rapidly out-competed by wild-type T4 in co-infection assays (Extended Data Fig. 6g).

Importantly, EM4 also bypasses PANGU*^Ecu^*-mediated defense (Extended Data Fig. 6h), implying that both PANGU types sense the same infection signal. Indeed, when we repeated the T4 evolution on PANGU*^Ecu^*-expressing cells, we obtained two additional escape mutants: one carries a single-nucleotide change in the *denA* ribosome-binding sequence; the other, a single-base insertion in *denA* that causes a frameshift (Extended Data Fig. 6i). Notably, both mutants also bypass PANGU*^Ff^*-mediated defense (Extended Data Fig. 6h). Collectively, our results show that T4 repeatedly mutates *denA* to escape PANGU defenses.

## EndoII is necessary but insufficient for induction of the Pag1A-Pag1B assembly

Next, we probed the Pag1A-Pag1B complex formation in T4 or EM4-infected cells through co-immunoprecipitation. N-terminally Flag-tagged Pag1B retained full immunity against T4 (Fig. 4d). After anti-Flag-precipitation of Pag1B, we probed for Pag1A with custom anti-Pag1A antibodies. As expected, Pag1A was efficiently co-immunoprecipitated from WT-T4-infected samples, whereas the signal was barely detectable after infection with EM4-T4, reflecting the markedly reduced complex formation in EM4-infected cells (Fig. 4d). Thus, T4 EndoII activity is required for the formation of a stable Pag1A-Pag1B complex.

Next, as a proxy for Pag1B activation, we assessed translation activity at different time points post-infection by puromycin incorporation^32^. During the infection with WT T4, the PANGU*^Ff^*system suppressed translation as early as 8 min (Fig. 4e), coinciding with their detected *in vivo* interaction (Fig. 4d). In contrast, EM4-infected cultures maintained puromycin labeling activity comparable to the vector control (Fig. 4e), further supporting the idea that EndoII deficiency prevents both complex formation and Pag1B-mediated translational shutdown.

EndoII alone, however, is insufficient for Pag1B activation. While expressing EndoII alone in PANGU*^Ff^* cells slowed the growth, presumably due to chromosomal fragmentation, it failed to elicit PANGU-dependent growth arrest or inhibition of translation (Extended Data Fig. 6e, f, j). Therefore, while EndoII initiates DNA cleavage, robust PANGU activation seemingly requires more extensive genome degradation (see below), which involves additional viral nucleases^17,30,31^.

## Pag1A binds to long DNA fragments

As a nucleoid-associated protein, Pag1A is expected to bind the *E. coli* chromosome *in vivo*. ChIP-seq showed that Pag1A associates with the *E. coli* chromosome in a seemingly sequence-nonspecific manner (Fig. 5a, Extended Data Fig. 7). A DNA-bound AlphaFold model of the Pag1A dimer reveals a central pore 22.6–26.8 Å across, wide enough to thread B-form DNA (∼20 Å) (Fig. 5b and Extended Data Fig. 2a). In the structure of the Pag1A₂-Pag1B heterotrimer, this pore collapses to only 4.1–9.5 Å (Fig. 3d and Extended Data Fig. 8a), indicating that Pag1B both occludes the entrance and constricts the channel, physically excluding DNA. Several basic residues (K/R) lining the pore are positioned to contact the phosphate backbone (Fig. 5b). Consistently, purified Pag1A retards a 1-kb dsDNA fragment (randomly selected from *E. coli* genome) in a gel-shift assay, whereas the triple-substituted Pag1A^KRK^ (K200A/R203A/K204A) variant fails to shift the probe (Fig. 5c and Extended Data Fig. 7b), confirming that Pag1A bind to DNA via these basic residues.

**Fig. 5.**
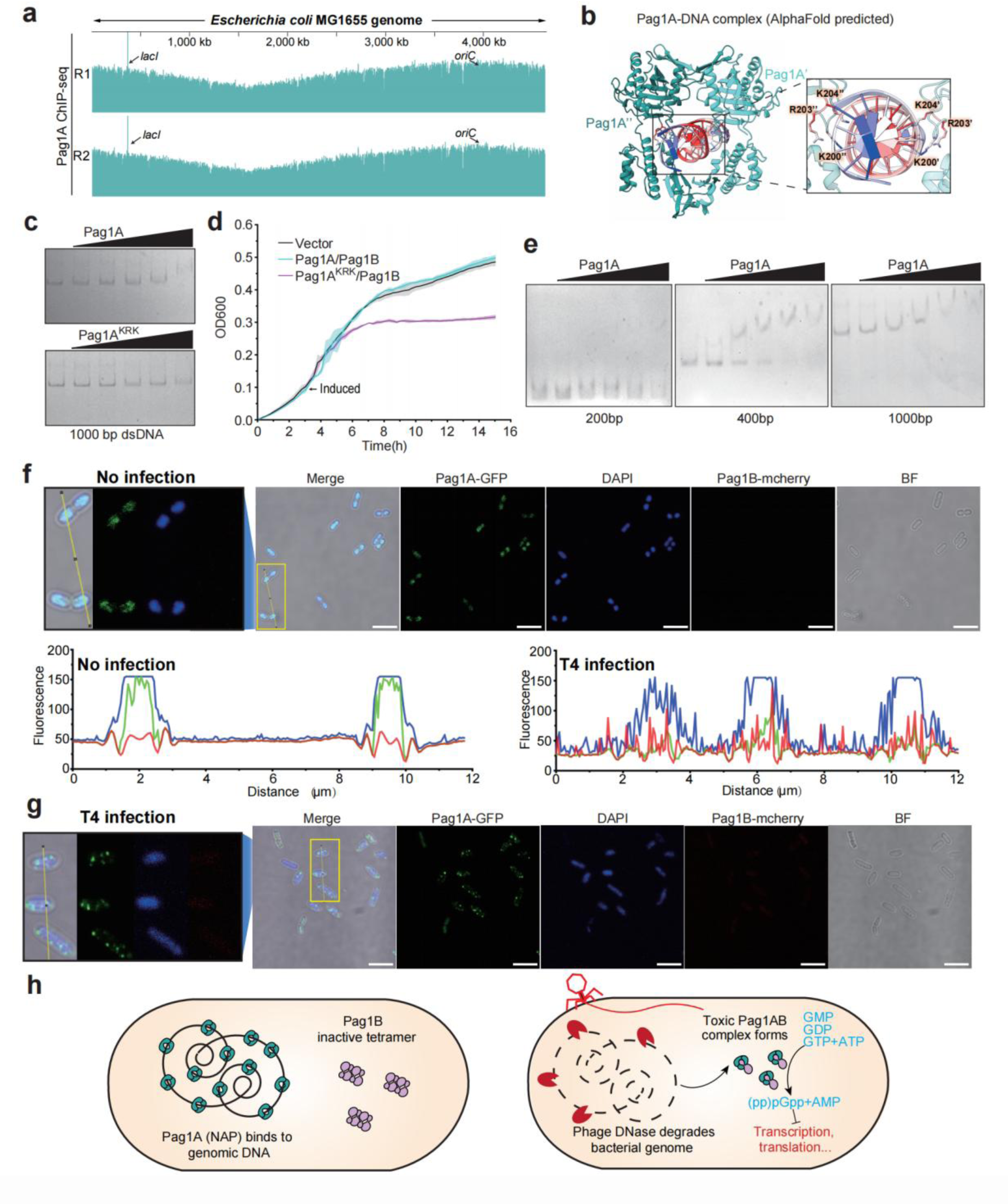
T4 infection induces the re-localization of Pag1A from the nucleoid to the cytoplasm. **a,** AlphaFold-predicted structure of DNA-bound Pag1A (see Extended Data Fig. 2a). DNA-contact residues K200, R203 and K204 are shown as sticks. **b,** EMSA of a 1-kb dsDNA probe incubated with wild-type (WT) Pag1A or the triple mutant Pag1A^KRK^ (K200A/R203A/K204A). FAM-labeled dsDNA was used at a fixed concentration of 10 nM, and Pag1A was tested at concentrations of 2, 4, 8, 16, or 32 μM. **c,** Growth curves of *E. coli* carrying empty vector (EV), PANGU*^Ff^*, or the variant system encoding Pag1A^KRK^. **d,** EMSA of Pag1A with dsDNA fragments of the indicated lengths. FAM-labeled dsDNA was used at a fixed concentration of 10 nM, and Pag1A was tested at concentrations of 4, 8, 16, 32, or 64 μM. **e,** ChIP-seq occupancy profile of Pag1A across the MG1655 genome. The *lacI* peak (left arrow) originates from the additional *lacI* copy on the pET28a backbone used for Pag1A expression. The right arrow indicates the location of replication ori. **f, g,** Fluorescence microscopy of PANGU*^Ff^*-expressing *E. coli* cells under uninfected (**f**) or T4-infected (**g**) conditions. Genomic DNA (DAPI stained), GFP-tagged Pag1A and mCherry-tagged Pag1B were visualized using the blue, green, and red channels, respectively. Merged images were analyzed using ImageJ’s Plot Profile tool to quantify signal correlation. The mCherry signal was weak because the chromophore matures slowly. Scale bar, 5 μm. **h,** Proposed model for the Type I PANGU immune system. In uninfected cells Pag1A is chromosome-bound and confined to the nucleoid region, whereas Pag1B forms an inactive tetramer within the cytoplasm. Upon T4 infection, phage nucleases degrade chromosomal DNA, releasing Pag1A into the cytoplasm where it disassembles the Pag1B tetramer. The resulting Pag1A₂-Pag1B heterotrimer converts ATP and GTP/GDP/GMP to AMP and alarmones, shutting down transcription, translation and other essential processes to abort infection.

Remarkably, co-expression of Pag1A^KRK^ with Pag1B strongly inhibited cell growth (Fig. 5d), confirming the idea that liberating Pag1A from chromosomal DNA is sufficient to unleash Pag1B toxicity. Interestingly, we further found that DNA length is critical, as the band of 200-bp fragments was not shifted by Pag1A even at high protein concentrations, whereas 400-bp and 1-kb DNA bands were readily shifted (Fig. 5e and Extended Data Fig. 7b). This explains why EndoII-mediated chromosome fragmentation (into ∼12-kb pieces) is insufficient to trigger PANGU defense, and more extensive genome degradation by other viral nucleases is required. Finally,

## Pag1A translocates out of nucleoid during T4 infection

The chromosome-binding capability of Pag1A and the assembly of Pag1A-Pag1B complex after viral induced genome degradation raised the possibility that Pag1A is released from the nucleoid to the cytoplasm (where Pag1B resides) during infection. To test this, we tagged Pag1A with green fluorescent protein (GFP) and Pag1B with mCherry to monitor their cellular localization using fluorescence microscopy. Pull-down assays showed that the fusion of GFP or mCherry does not impair Pag1A-Pag1B interaction (Extended Data Fig. 8b). Next, nucleoid DNA was visualized with DAPI staining. In uninfected cells, Pag1A-GFP signal overlapped precisely with DAPI-stained nucleoid (Fig. 5f), confirming its tight association with the nucleoid. Conversely, in T4-infected cells, Pag1A-GFP diffused into the cytoplasm, forming discrete foci, and was no longer co-localized with the DAPI signal (Fig. 5g), indicating its release from the nucleoid. In contrast, Pag1B-mCherry remained evenly distributed throughout the cytoplasm under both conditions (Fig. 5f, g).

In summary, the collective data presented here support the following model for phage sensing and restriction by Type-I PANGU (Fig. 5h). Prior to infection, NAP protein Pag1A is sequestered on nucleoid DNA and is physically separated from the enzymatically inactive Pag1B homotetramer. Early in T4 infection, phage-encoded nucleases, initiated by EndoII, degrade the host chromosome, thereby liberating Pag1A. The freed Pag1A migrates to the cytoplasm, disassembles the Pag1B tetramer, and assembles the active Pag1A-Pag1B heterotrimer. Activated Pag1A-Pag1B, in turn, depletes ATP and GTP while synthesizing alarmones, thus blocking essential cellular processes and aborting phage propagation.

Generality of the Pag1A-mediated sensing mechanism is supported by the observation of a similar relocation upon triggering of the Type-II PANGU*^Ecu^*system (Extended Data Fig. 8c, d). GFP-tagged Pag2A co-localized with the nucleoid in uninfected cells, but redistributed to a diffuse cytoplasmic pattern (without distinct foci) upon T4 infection. Thus, phage-induced re-localization of the core NAP sensor appears to be a shared feature of Type I and Type II PANGU systems; how Pag2A engages and activates its cognate effector Pag2B remains to be determined.

## PANGU-like defenses employ diverse effectors

To explore the diversity of PANGU-like defense systems, we first constructed a Hidden Markov Model (HMM) profile to search for YejK-like NAP proteins in NCBI GenBank, yielding a dataset of 1,638,667 NAPs (Fig. 6a). Notably, 47.1% (771,801) of the detected NAPs reside in putative operons, defined as adjacent genes separated by less than 40 bp, implying widespread partnering with non-NAP proteins. Because prokaryotic defense genes frequently cluster into “defense islands” ^33,34^, we retained only non-redundant NAP operons located within 10 kb of known immune loci, yielding 2,092 non-redundant defense-linked operons (Fig. 6a). This collection constituted approximately 8.22% of the 25,501 non-redundant NAP-encoding operons, versus 4.20 % (843 out of 20,076) for standalone NAPs, a difference that is significant across major phyla (Extended Data Fig. 9a). It appears that operon-encoded NAPs are more likely to co-localize with defense systems.

**Fig. 6.**
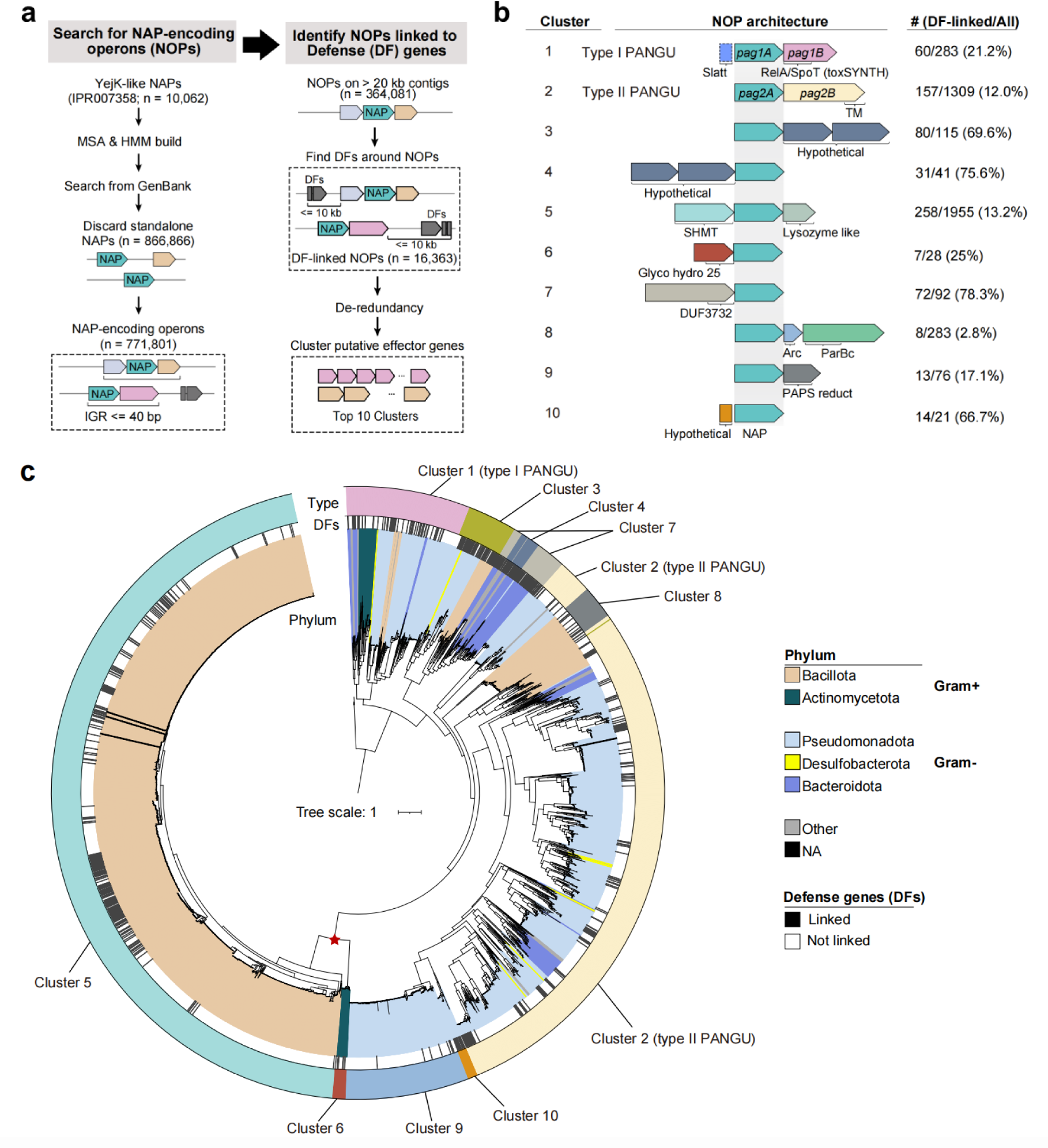
Systematic search and phylogenetic analysis of PANGU-like defense systems. **a,** Workflow for identifying defense-related NAP-encoding operons (NOPs). A bioinformatic search for NOPs were first performed, followed by analyses of their genetic linkage to known defense genes (DF, dark grey). Candidate NOPs retained after each filter (discarding standalone NAPs, removing NOPs on short contigs, or depleting redundant sequences) are highlighted in dashed boxes. Putative effectors encoded by the final dataset of defense-related NOPs were clustered by sequence similarity, with the top 10 largest clusters summarized in panel B. **b,** Top 10 clusters of defense-related NOPs (putative PANGU-like defense systems). The dashed box indicates the presence of the SLATT gene in a subset of Cluster 1 NOPs. The number (#) of candidates in each cluster (DF-linked), and their occurrence in the dataset of all NOPs are indicated, as well as the percentage of DF-linked candidates. **c,** Phylogenetic relationships of NAPs from top 10 NOP clusters. From the inner to outer circles: (1) phylum-level classification of NAP-containing strains (color-coded; Gram-positive = Gram+, Gram-negative = Gram−); (2) genetic linkage to known defense genes (DFs); (3) Cluster designation (Cluster1-Cluster10). The red star marks a shared ancestral node between Cluster5 and Cluster6.

Hierarchical clustering of the neighboring genes produced dozens of NAP-centered operon types, with the ten largest designated Cluster 1-10 (Fig. 6b). Cluster1 and Cluster2 correspond to Type I and Type II PANGU, respectively. Several Cluster1 operons additionally encode a SLATT-domain protein alongside the GRel toxin Pag1B. Cluster3 and Cluster4 carry two conserved hypothetical proteins, located downstream and upstream of the core NAP gene, respectively. Cluster5 and Cluster6 couple the NAP to distinct lysozyme families, suggesting shared peptidoglycan-targeting effector strategies. Cluster7 (encoding a DUF3732-domain protein) and Cluster10 (encoding a small hypothetical protein) operons co-localize with other defenses at frequencies of 78 % and 67 %, respectively, indicating their high potential in anti-phage defense. Cluster8 pairs the NAP with a PAPS reductase, and Cluster9 adds two additional proteins featuring Arc and ParBc domains. These findings highlight the modular nature of PANGU-related defenses, which likely enhances their functional versatility and facilitates their evolutionary adaptation to diverse hosts facing different phage threats.

## Wide distribution and polyphyletic origin of PANGU-related defenses

A maximum-likelihood tree of the core NAP proteins reveals distinct clade separation among the ten clusters (Fig. 6c). Phylogenetic analysis including standalone NAPs showed that they occupied ancestral branches in each clade, whereas PANGU-related NAPs formed tight terminal clusters with extremely short branch lengths, creating characteristic “plateau” structures (Extended Data Fig. 9b). This topology suggests a polyphyletic origin for PANGU-related defenses, and their functional stabilization and vertical inheritance following system establishment.

Taxonomic profiling places the majority of PANGU systems in Bacillota (Gram-positive) and Pseudomonadota (Gram-negative), with sporadic representation in Bacteroidota, Actinomycetota, Desulfobacterota, and others (Fig. 6c). Cluster1 (Type I) PANGU systems distribute across over five phyla, indicating their high compatibility. Besides, their leaf nodes located in clades close to the root suggest their early emergence during evolution. Similarly, the interphylum distribution of Cluster2 (type II PANGU systems), Cluster3, Cluster4, and Cluster7 implies their spread through horizontal transfer events. By contrast, Cluster5 and Cluster8 are confined to Bacillota, and Cluster9 and Cluster10 to Pseudomonadota, suggesting that these clusters may have emerged more recently or undergone limited cross-phylum horizontal transfer. Notably, Cluster5 and Cluster6 candidates share a common ancestral node, indicating a shared evolutionary origin. In the global NAP phylogeny, standalone NAPs show more inter-phylum jumping than operon-associated ones (Extended Data Fig. 9b), suggesting that once combined with a defensive effector, the locus experience reduced horizontal transfer, likely due to functional constraints or selective pressures that stabilize their genomic context within host defense mechanisms.

Collectively, our findings indicate that PANGU-related defenses likely originated from ancestral NAPs that acquired diverse effector partners during horizontal transfers, supporting a polyphyletic origin and wide distribution. Their ability to augment host adaptability under diverse selective pressures likely drove their retention and vertical transmission, leading to the current widespread and functionally diverse repertoire of PANGU-related defenses.

## Discussion

Phages frequently encode nucleases to degrade the bacterial genome DNA, yielding substrates for viral DNA replication^17,30,31^. The Hachiman heterodimeric complex HamAB has recently been reported to sense DNA damages, and upon this detection, helicase activity of HamB unleashes the nuclease activity of HamA, leading to nonspecific DNA cleavage within the cell^35^. Our study reveals a novel group of abortive anti-phage systems that use YejK-like nucleoid-associated proteins (NAPs) to monitor the integrity of the host genome. Upon the degradation of the bacterial genome by viral nucleases NAPs are released from the nucleoid and translocate to the cytoplasm (or possibly the cell membrane), where they bind to and activate toxic immune effectors. This activation halts critical cellular processes, such as transcription and translation, thereby preventing phage propagation. Such defense systems termed PANGU force phages to an evolutionary trade-off between taking over host nucleotide resources and escaping host immune surveillance. Notably, other coliphages (e.g., T1 and T5) that degrade host genome during infection^36^ were found to be resistant to the two PANGU systems we investigated (Extended Data Fig. 10). This suggests either that the extent of genome degradation by these phages is insufficient to trigger these PANGU systems, or that they encode potential anti-PANGU factors, which warrants further investigation. Afterall, PANGU systems highlight a unique immunity controlling strategy based on spatial separation of sensor and effector modules, combined with a distinct activation mechanism by phage infections that relies on the cellular redistribution of the sensor module.

Our study reveals that PANGU-like defense systems are widespread across both Gram-negative and positive phyla, with YejK-like proteins serving as key components, thus highlighting their broader, previously unrecognized role in bacterial immunity. Importantly, bioinformatic analyses indicate that nearly half of all identified YejK-like NAPs are organized into operons with one or more additional genes, suggesting that these NAPs often collaborate with diverse effector proteins to mediate functions like immunity. While we have initially defined 10 major clusters of PANGU-like systems based on their genetic linkage to known defense systems, the diversity of these types implies that additional types remain to be uncovered, particularly those outside defense islands. Further investigation into their immune mechanisms is expected to uncover further diversity in NAP-based defense strategies. It will also be interesting to investigate the coevolution between the NAP sensor and different effectors, as AlphaFold ipTM scores and Pearson correlations indicate that Pag1A–Pag1B pairs have co-evolved with each other rather than with their host genomes (Extended Data Fig. 11).

Type I PANGU employs a toxSAS GRel toxin, Pag1B, as an anti-phage effector that converts ATP and GTP to AMP and pppGpp to shut down essential cellular pathways and abort infection. This effector is related to but is fundamentally different from anti-phage toxSAS CapRel, a fused toxin-antitoxin that senses phage proteins and, upon interaction, pyrophosphorylates tRNAs to induce an abortive response^12,24^. While translation-targeting CapRel is auto-inhibited by intramolecular contacts between its toxin and antitoxin domains and is triggered when phage proteins disrupt this interface^12^, full activation of the metabolism-targeting GRel/Pag1B strictly requires allosteric activation by Pag1A. These contrasts underscore the mechanistic diversity of toxSAS-family toxins and highlight their expanding repertoire in bacterial immunity.

In summary, PANGU defines a distinct, highly modular, and widely distributed class of anti-phage defenses, and illustrates a unique infection sensing mechanism based on the re-localization of a nucleoid-associated sensor. Our findings also highlight the previously unrecognized, yet critical, role of abundant bacterial NAPs as sentinels in the immune landscape.

## Methods

### Bacterial strains and bacteriophages

*E. coli* DH5α was used for plasmid construction, and *E. coli* BL21 (DE3) strain was used for protein expression. *E. coli* MG1655 and BW25113 were used as the host *E. coli* strain for each phage. All bacterial cultures were cultured at 37°C in Lysogeny Broth (LB) medium composed of 10 g/L tryptone, 5 g/L yeast extract, and 10 g/L NaCl. Solid agar plates were prepared by supplementing LB medium with 15 g/L agar, while liquid cultures were grown with shaking at 160-200 rpm. Phages were propagated in liquid LB medium, followed by centrifugation and filtration through a 0.22-μm filter to remove cellular debris. Phage titers were determined via the small-drop plaque assay method, which allows for precise quantification of infectious phage particles.

### Plasmid construction and transformation

The synthetic genes, primers and plasmids used in this study are detailed in Supplementary Table. The synthetic PANGU locus sequence was generated by BGI Write (Beijing, China). Double-stranded DNA fragments were amplified using Phanta Super-Fidelity DNA Polymerase (Vazyme Biotech, P505-d2). Subsequently, these fragments were assembled into a predigested vector using the Hieff Clone® Plus One Step Cloning Kit (Yeasen Biotech, 10911ES50) via the Gibson assembly strategy. Constructed plasmids were verified by Sanger DNA sequencing (Tsingke, Beijing, China). PANGU*^Ec^* was cloned into the pBR322-derivative^37^ downstream of the Ptet constitutive promoter. NA37 and gRel were ordered as gBlocks to Integrated DNA Technologies, with the native Shine-Dalgarno sequence and flanking regions for Gibson assembly (GA). VHP1524 plasmids were verified by Sanger DNA sequencing (LGC Genomics GmbH, Berlin, Germany). For bacterial transformation, *E. coli* DH5α and *E. coli* BL21(DE3) competent cells (prepared using standard heat-shock methods) were used for cloning and protein expression, respectively. *E. coli* MG1655 competent cells were generated using the Ultra-Competent Cell Prep Kit (Sangon Biotech, B529303-0200). For transformation, 200 ng of plasmid DNA was added to 100 μL of competent cells and incubated on ice for 30 minutes. Transfer the cell-DNA mixture to a preheated (42°C) water bath and incubate for 45 seconds. Immediately return to ice for 2–3 min to allow DNA uptake. Following a 1-hour recovery culture at 37°C in 900 μL of LB medium, cultures were serially diluted (10-fold) and plated onto LB agar plates containing appropriate antibiotics, and successful transformants were verified by colony PCR.

### Plaque assays

Plaque assays were performed using a double agar overlay protocol. Single colonies of *E. coli* MG1655 harboring a pET28a-PANGU or empty plasmid was grown overnight at 37°C in LB medium. For overlay preparation, 100 μL of a saturated bacterial culture was mixed with molten LB agar (0.7% w/v) supplemented with 50 μg/mL kanamycin and isopropyl-β-D-thiogalactopyranoside (IPTG) at concentrations of 0.1 mM (for PANGU*^Ecu^*), or 1 mM (for PANGU*^Ff^*). The top agar mixture was poured onto a prepared LB agar bottom layer (1.5% w/v agar) and allowed to solidify at room temperature for 15 minutes. Phages were serially diluted (10-fold) in LB medium, and 2 μL of each dilution were spotted onto the overlay and air-dried. Plates were incubated at 37°C for 12–16 hours to allow plaque formation. After incubation, plates were scanned using a standard flatbed scanner, and plaque-forming units (PFU) were enumerated by counting distinct plaques, keeping note of changes in plaque size relative to a negative control. The efficiency of plating (EOP) was calculated as the ratio of PFU in experimental conditions to PFU in the negative control (mean PFU experimental / mean PFU negative control). Overnight cultures of *E. coli* BW25113 carrying either VHp1524 or the empty vector VHp1423 were diluted in top agar (0.5%) to a final concentration of 0.075 OD600 cells/mL and poured onto LB agar plates (1.5%). Top agar layer was left at room temperature for 15 minutes to solidify. 10-fold serial dilutions of Bas34 were prepared with SM buffer and 2.5 μL of each dilution were spotted onto the top agar. Plates ere incubated at 37°C and PFU were counted after 6 and 24 hours to determine the EOP. All assays were performed in biological triplicate to ensure statistical robustness, and data visualization was conducted using GraphPad Prism.

### Infection dynamics in liquid culture

Single colonies of *E. coli* MG1655 harboring the pET28a-PANGU or empty plasmid was grown overnight at 37°C in LB medium. The cultures were diluted 1:100 in fresh LB medium supplemented with 50 μg/mL kanamycin and IPTG at concentrations identical to those used in the plaque assays. The cultures were incubated at 37°C with shaking at 200 rpm, then 180 μL early-logarithmic phase cultures were transferred into wells of a 96-well plate containing 20 μL of phage diluent to achieve final multiplicity of infection (MOI) of 0.5 or 10 for phage, while 20 μL of LB medium was added to uninfected control wells. Infections were performed in triplicate, using cultures derived from three independent bacterial colonies to ensure experimental reproducibility. Cell viability and lysis kinetics were monitored by periodically measuring OD600 every 15 minutes using a real-time plate reader (Stratus 600; Cerillo, Charlottesville, VA, USA). The culture plate was continuously shaken at 37°C during measurement to maintain homogeneous cell distribution. Data were collected and analyzed using GraphPad Prism.

### Structure and domain analysis

Structures were predicted using AlphaFold3 Server^38^ (https://golgi.sandbox.google.com/). Domain analysis was performed using HHpred (https://toolkit.tuebingen.mpg.de/tools/hhpred).

### Protein purification

The protein purification protocol was optimized for enhanced solubility and resolution. Pag1A and Pag2A were C-terminally tagged with 6xHis-Flag, while DMK (Pag2B) was N-terminally tagged with 6xHis. For Pag1B, an N-terminal 6xHis-SUMO tag was introduced to improve solubility. The recombinant plasmids were transformed into *E. coli* BL21(DE3). Expression was induced with 0.5 mM IPTG at 22°C for 16 hours. Cell pellets were resuspended in binding buffer (20 mM Tris-HCl, pH 7.5, 500 mM NaCl, 20 mM imidazole) and lysed by sonication. The cleared lysate was loaded onto a His-Trap HP column (GE Healthcare) pre-equilibrated with binding buffer. After washing with wash buffer (binding buffer containing 50 mM imidazole), the protein was eluted using a linear gradient of 500 mM imidazole in the same buffer. Purified proteins were further separated by size-exclusion chromatography (SEC) on a Superdex 200 10/300 GL column (GE Healthcare) using storage buffer (20 mM HEPES, pH 7.5, 150 mM NaCl, 5% glycerin) as the mobile phase. Fractions containing the target protein were pooled, concentrated. The fusion protein was then digested with SUMO Protease (Beyotime Biotech, P2312S) at 4 °C overnight to remove SUMO tag. The concentration of the recombinant protein was determined using a BCA protein assay kit (Thermo Scientific, 23227) and verified by SDS-PAGE under reducing conditions, with Coomassie Brilliant Blue G250 staining (50% methanol, 5% acetic acid, 0.1% Coomassie) followed by destaining with water.

### *In vitro* Flag pull-down assay

The purified Flag-Pag1A fusion protein was incubated with Pag1B at a 1:2 molar ratio using anti-FLAG M2 magnetic beads (Sigma, M8823) under gentle rotation at 4°C for 3 hours to facilitate protein-protein interaction capture. A parallel negative control was performed by incubating Pag1B alone with magnetic beads under identical conditions. Following incubation, the supernatant was carefully aspirated, and the beads were washed four times with ice-cold wash buffer (20 mM HEPES, pH 7.5, 150 mM NaCl, 1% Triton X-100) to remove non-specifically bound proteins. The bound complex was then eluted using 3X Flag peptide-containing elution buffer (20 mM HEPES, pH 7.5, 150 mM NaCl, 1% Triton X-100, 0.4 mg/mL 3X Flag Peptide), which specifically displaces the Flag-tagged protein through competitive binding. The eluted samples were subsequently analyzed by SDS-PAGE using a 12% separating gel and Coomassie Brilliant Blue staining to verify protein integrity and purity.

### BLI assay

BLI assays were performed using the Octet RED96 system (Sartorius) to determine the binding kinetics of Pag1A/Pag2A with DMK(Pag1B)/DMK(Pag2B). Recombinant receptors were biotinylated with EZ-Link NHS-PEG12-biotin (Thermo Scientific, 21312) and subsequently immobilized onto streptavidin (SA) biosensors (ForteBio, 1912052) at a concentration of 10 μg/mL. All assays were conducted in 200 μL of assay buffer (20 mM HEPES, pH 7.5, 150 mM NaCl, and 0.02% Tween-20) within 96-well black flat-bottom plates maintained at 37°C with orbital shaking at 1,000 rpm. During the association phase, biosensors were exposed to DMK(Pag1B)/DMK(Pag2B) at varying concentrations, followed by a dissociation phase in assay buffer. Kinetic data were analyzed using ForteBio Data Analysis Software and data visualization with GraphPad Prism.

### Cryo-EM specimen preparation and data collection

For the preparation of cryo-EM specimens of Pag1A-Pag1B complex, 3.5 μL sample was applied to glow-discharged holey carbon copper grids (Quantifoil R 1.2/1.3, 300 mesh). After blotting for 2 seconds, specimens were vitrified using a Vitrobot Mark IV (Thermo Fisher Scientific). Data were acquired using a CRYO ARM 300 electron microscope (JEOL) at 300 kV, with a K3 direct electron detector (Gatan). Images were captured automatically by Serial-EM software in super-resolution mode, setting the physical pixel size at 0.95 Å and defocus range between -1 and -2.5 μm. Each image stack was divided into 40 frames, with a total electron dose of 50 e/Å2.

### Cryo-EM data processing and model building

Data processing was consistent across all data sets using cryoSPARC v.4.4.1. Procedures included patch motion correction, patch CTF estimation, and particle picking. Three rounds of 2D classification were applied after the blob picking and particle extraction. Finally, only 226,103 particles were selected for ab-initial reconstruction and hetero-refinement. Non-uniform refinement was then applied to achieve the density map at 2.67 Å. Local refinement was used to further optimize the map and local resolution estimation was carried out. The final resolutions achieved were 2.64 Å.

The initial structural model was generated by CryFold. The complex structure was then manually fixed in COOT and refined using the Phenix.real_space_refine module in PHENIX.

### Cell-free translation

The PURExpress *in vitro* protein synthesis kit (NEB, E6800) was employed following the manufacturer’s protocol. Each reaction mixture was supplemented with 0.8 U/μL RNase Inhibitor Murine (NEB, M0314S) to minimize RNA degradation. Purified Pag1A/Pag1B protein was used at final concentrations of 62.5, 125, 250, and 500 nM, alongside a constant concentration of 10 ng/μl DHFR template plasmid. Mock control reactions were conducted by substituting Pag1A/Pag1B protein with an equal volume of storage buffer. After a 1-hour incubation at 37 °C, reaction aliquots (5 μL) were mixed with an equal volume of 4 × sample buffer (250 mM Tris-HCl, pH 6.8, 10% SDS, 0.5% bromophenol blue, 50% glycerol, and 5% β-mercaptoethanol), boiled for 5 min at 98 °C, and resolved by 12% SDS-PAGE. After staining with Coomassie Brilliant Blue, the quantity of DHFR expression products was quantified.

### Enzymatic assays: alarmone synthesis by Pag1A and Pag1B

Reaction mixtures containing 2 mM NTP or ATP and GTP/GDP/GMP /dNTP/ITP, 16 μM Pag1A or/and 16 μM Pag1B in a 50 μL reaction buffer (20 mM HEPES, pH 7.5, 150 mM NaCl, 5 mM MgCl₂, 1 mM DTT, 5% glycerol) were incubated at 37 °C for 2 hours. The reactions were quenched with methanol. Precipitated protein was removed by centrifugation, and the resulting supernatant was subjected to LC-MS analysis. Separation was performed on a Shimadzu UPLC system coupled to a QTOF mass spectrometer using a C18 analytical column (AQ-C18, 250 × 4.6 mm, 5 μm particle size; Welch Technologies). The mobile phase consisted of 5 mM ammonium formate in water (pH adjusted to 8.5 with ammonium hydroxide; solvent A) and acetonitrile (solvent B). The following gradient was applied at a flow rate of 0.6 mL/min: 0–7 min, 1% B; 7–10 min, 1–10% B; 10–15 min, 10–80% B; 15–18 min, 80% B; 18–20 min, 80–1% B.

### Enzymatic assays: tRNA pyrophosphorylation by Pag1A and Pag1B

The reaction mixture containing 5 μM total deacylated tRNA preparation from *E. coli* (Sigma-Aldrich, R1753), 500 μM [γ³²P]ATP, and 500 nM Pag1A or/and Pag1B in reaction buffer (20 mM HEPES, pH 7.5, 150 mM NaCl, 5 mM MgCl₂, 1 mM DTT, 5% glycerol) was incubated at 37 °C for 1 hour. To detect phosphorylated tRNA, the sample was mixed with an equal volume of 2× RNA loading buffer (98% formamide, 10 mM EDTA, 0.3% bromophenol blue, and 0.3% xylene cyanol), heated at 90 °C for 5 minutes, and resolved by urea-PAGE (8% polyacrylamide, 8 M urea) in 1× TBE buffer. Following electrophoresis, the gel was dried and exposed to X-ray film in a light-tight cassette at -70 °C. Radioactive signal was detected by autoradiography. In parallel, tRNA was visualized using M5 GelRed Plus (Mei5 Biotech, MF079-plus-01) under UV transillumination.

### *In vitro* transcription assay

*In vitro* transcription reaction was performed using a DNA template containing a 500-bp sequence under the control of the Tac promoter. The 50-μL *in vitro* transcription reaction was prepared in transcription buffer (40 mM Tris-HCl, pH 7.5, 150 mM KCl, 10 mM MgCl₂, 1 mM DTT, 0.01% Triton® X-100) and supplemented with 2 units of RNAP holoenzyme (NEB, M0551S) and 50 nM DNA template, in the presence or absence of 16 μM Pag1A and Pag1B. RNA synthesis was initiated by adding NTPs at a final concentration of 0.5 mM each, followed by incubation at 37°C for 1 hour. To visualize the RNA products, the sample was mixed with an equal volume of 2× RNA loading buffer (98% formamide, 10 mM EDTA, 0.3% bromophenol blue, and 0.3% xylene cyanol). The mixture was heated at 90 °C for 5 minutes and immediately chilled on ice. RNA transcripts were separated by electrophoresis on a 1% agarose gel in 1× TAE buffer and visualized using M5 GelRed Plus (Mei5 Biotech, MF079-plus-01) under UV transillumination.

### Size-exclusion chromatography (SEC) analysis

To investigate the oligomeric states, purified Pag1A and Pag1B were resolved by size-exclusion chromatography (SEC) on a Cytiva HiLoad 16/600 Superdex 200 pg column. Fractions (3 mL each) were collected and assayed by OD280. For the Pag1A and Pag1B complex formation assay, purified Pag1A and Pag1B were incubated in an equimolar ratio for 30 min at 4 °C prior to SEC fractionation. Co-eluting proteins were identified by SDS–PAGE, and gels were stained with Coomassie Brilliant Blue R-250 for protein visualization.

### SEC-MALS

Molecular Weight Determination by SEC-MALS was performed on a Vanquish Core HPLC system (Thermo Fisher Scientific, Waltham, MA, USA) connected to a DAWN multi-angle light scattering detector and an Optilab differential refractometer (Wyatt Technology, Santa Barbara, CA, USA). The system was equilibrated with buffer containing 150 mM NaCl, 50 mM Tris-HCl (pH 7.5), and 0.02 mM TCEP. Purified samples (50 μL each) of Pag1A, Pag1B, or Pag1A-Pag1B complexes were injected and eluted at a flow rate of 0.75 mL/min at room temperature. Light scattering and refractive index signals were recorded and analyzed using ASTRA software (Wyatt Technology). The light scattering detector was calibrated using a calibration constant of 5.7351 × 10⁻⁵ cm⁻¹ V⁻¹. The refractive index of the solvent was set to 1.329, and the refractive index increment (dn/dc) was defined as 0.185 mL/g.

### Native electrophoresis analysis

To analyze the formation of Pag1A and Pag1B complexes, 10 μM of each protein was incubated in 10 μL of binding buffer (20 mM HEPES, pH 7.5, 150 mM NaCl, 5% glycerol) at 37 °C for 30 minutes. In the gradient dissociation experiment of Pag1A from Pag1B, the concentration of B was maintained at 10 μM, while the concentration of A was serially diluted from 10 to 5, 2.5, and 1.25 μM. For Native PAGE, the samples were mixed with 0.5% Coomassie Brilliant Blue G-250 and NativePAGE sample buffer (Invitrogen, BN2003), then resolved on precast NativePAGE Bis-Tris Mini Protein Gels (Invitrogen, BN1001BOX). Electrophoresis was performed at 150 V for 115 minutes using NativePAGE running buffer (Invitrogen, BN2007). Following electrophoresis, the gels were stained with Coomassie Brilliant Blue G-250 staining solution (0.1% Coomassie Brilliant Blue G-250, 50% methanol, 5% acetic acid) for 30 minutes and destained in deionized water

### Isolation of phage escape mutants against PANGU

The phage evolution experiment was conducted as described previously^28^. In brief, independent populations were evolved in a 96-well plate containing a sensitive host *E. coli* MG1655 pET28a (empty vector) and a resistant host PANGU defense system-expressing *E. coli* MG1655 pET28a-PANGU. One control population was evolved with only the sensitive host. Overnight bacterial cultures were back-diluted to OD600 = 0.1 in LB medium and 180 μL were seeded into each well. Cells were infected with 20 μL tenfold serial dilutions of T4 phage with MOI from 10 to 0.0001, with one well uninfected to monitor for contamination. Plates were incubated at 37 °C for 8 h in a plate shaker at 1,000 rpm. Following each evolutionary cycle, lysates from cleared wells were pooled and centrifuged (5,000 × g, 10 min) to remove cellular debris, and the supernatant lysates were transferred to a 96 deep-well block with 40 μl chloroform added to prevent bacterial growth. Lysates were spotted onto both sensitive and resistant hosts to check the defense phenotype. This selection regime continued iteratively until all experimental populations demonstrated the ability to overcome the PANGU defense system. Evolved clones were isolated by plating to single plaques on lawns of resistant host, and control clones from the control population were isolated on a lawn of the sensitive host. One representative clone from each population were propagated using the corresponding host followed by whole-genome sequencing.

### Phage DNA extraction and Illumina sequencing

The genomic DNA of phage was extracted using the Lambda Phage Genomic DNA Kit (Zoman Biotech, ZP317) following the manufacturer’s protocol. In brief, high-titre phage lysates (>106 PFU/μL) were treated with DNase I (0.001 U/μL) and RNase A (0.05 mg/mL) at 37 °C for 30 min. Precipitate phage using a phage precipitation solution, followed by lysing the phage with a lysis solution. Lysates were then incubated with proteinase K at 56 °C for 1-hour to disrupt capsids and release phage DNA. Subsequently, DNA was extracted using the column-based method, and then eluted with 50 μl TE buffer (10 mM Tris-HCl, pH 8.0, 0.1 mM EDTA). Concentrations of extracted DNA were measured by NanoDrop (Thermo Fisher Scientific).

Phage genome resequencing was performed by GENEWIZ (Suzhou, China). Genomic DNA is fragmented, followed by end repair and 3’ adenylation to generate A-overhangs for adapter ligation. Adapters with P5/P7 sequences and dual-index barcodes are ligated to fragments, which undergo size selection using magnetic beads. Limited-cycle PCR enriches adapter-ligated fragments and incorporates sample-specific indices, with library quantification and fragment size verification. Denatured DNA is immobilized on a flow cell for bridge amplification. Paired-end sequencing (2×150 bp) is performed on the Illumina HiSeq/NovaSeq platform. Raw data undergoes adapter trimming, quality filtering, and alignment to the reference genome. PCR duplicates are removed, and coverage depth, uniformity, SNPs, and InDels are calculated for genome-wide variant detection.

### Nascent protein synthesis determined by puromycin incorporation

Puromycin incorporation was employed to quantify nascent protein synthesis^32^. Single colonies of *E. coli* MG1655 harboring pET28a-PANGU*^Ff^* or empty plasmid were cultured to the logarithmic phase (OD600 = 0.5) and subsequently infected with T4 or EM4 at a MOI of 10. Puromycin (Sigma, P8833) was added at sequential time points post-infection to achieve a final concentration of 10 μg/mL, followed by incubation at 37°C for 10 minutes to allow its incorporation into nascent polypeptides. Cells were harvested by centrifugation, washed once with precooled PBS, and lysed using RIPA Complete Lysis Buffer (Beyotime Biotech, P0013). Cell extracts were quantified by BCA protein assay kit and 5 μg of total protein was subjected to Western blot analysis. Blots were probed with monoclonal anti-puromycin antibody (1:5000 dilution, Sigma-Aldrich, ZMS1016) followed by HRP-conjugated anti-mouse secondary antibody (1:5000 dilution, Epizyme Biotech, LF101) to detect puromycin-labeled nascent peptides.

### Co-Immunoprecipitation

The interaction of Pag1A with Pag1B *in vivo* was detected by Co-Immunoprecipitation (Co-IP) using anti-FLAG M2 magnetic beads (Sigma, M8823). Single colonies of *E. coli* MG1655 harboring pET28a-derived plasmids expressing Pag1A and Flag-tagged Pag1B were cultured overnight in LB medium supplemented with 50 μg/mL kanamycin. Cultures were diluted 1:100 into fresh LB medium supplemented with 50 μg/mL kanamycin and 1 mM IPTG and incubated at 37 °C with shaking at 200 rpm until an OD600 of 0.6 was reached. At this experimental time point, formaldehyde crosslinking was initiated for the untreated control group, while the other two group cultures were further incubated with T4 phage or EM4 phage for 20 minutes prior to crosslinking. Formaldehyde was directly added to the culture medium to a final concentration of 1% (v/v) to covalently crosslink proteins and DNA, thereby stabilizing the Pag1A-DNA interaction during subsequent cell lysis and immunoprecipitation procedures. This formaldehyde-based crosslinking protocol effectively preserved Pag1A-DNA interactions under stringent experimental conditions, thereby minimizing the background noise of the interaction between Pag1A and Pag1B. Cells were lysed using RIPA Complete Lysis Buffer containing protease inhibitors (Roche, 04693132001), followed by centrifugation at 12,000 rpm for 10 minutes at 4°C. The supernatant was incubated with prewashed anti-FLAG M2 magnetic beads for 3 hours at 4°C with gentle shaking. The beads were washed five times with ice-cold lysis buffer and eluted using elution buffer (20 mM HEPES, pH 7.5, 150 mM NaCl, 1% Triton X-100, 0.4 mg/mL 3X Flag Peptide). Eluted proteins were resolved on a 12% SDS-PAGE and analyzed by Western blot using anti-FLAG, anti-HA, and anti-Pag1A antibodies. The experiment was performed in biological triplicate to ensure reproducibility.

### Western blotting

Western blot analysis was conducted to investigate the expression and interaction of Pag1A and Pag1B in both cell lysates and Co-IP samples derived from tagged Pag1A/Pag1B expressing cells. Protein samples were resolved by 12% SDS-PAGE and subsequently transferred onto a nitrocellulose membrane using a wet transfer system. Blots were blocked with 5% non-fat milk in TBST for 1-hour at room temperature and then incubated with primary antibodies targeting Flag (1:5000, Epizyme Biotech, LF303), HA (1:5000, Epizyme Biotech, LF313), and Pag1A (1:5000) sequentially overnight at 4°C. After thorough washing with TBST (4×5 min), HRP-conjugated secondary antibodies (1:5000) were applied for 1-hour at room temperature. Chemiluminescence signals were visualized using the Omni-ECL™ Femto Light detection kit (Epizyme Biotech, SQ201) and recorded on a Tanon-5200 Multi imaging system (Tanon Science & Technology).

### ChIP-seq

Single colonies of *E. coli* MG1655 harboring pET28a-derived plasmids expressing FLAG-tagged Pag1A and Pag1B were cultured overnight in LB medium supplemented with 50 μg/mL kanamycin. The cultures were diluted 1:100 into fresh LB medium containing 50 μg/mL kanamycin and 1 mM IPTG, followed by incubation at 37 °C with shaking at 200 rpm until the OD600 reached 0.6. At this time point, mix the sample with a 1% formaldehyde solution and crosslink it at 37°C for 15 minutes. Then, add 2.5 M glycine to a final concentration of 125 mM and shake at 37°C for 10 minutes to stop the crosslinking process. Cells were then collected by centrifugation at 9000 × g for 5 minutes. The cell pellets were washed two times with PBS buffer, flash-frozen in liquid nitrogen, and stored at −80 °C. For chromatin immunoprecipitation (ChIP), cell lysates were prepared by resuspending the pellets in lysis buffer (50 mM HEPES, pH 7.5, 150 mM NaCl, 1 mM EDTA, 0.1% sodium deoxycholate, and 1% Triton X-100) supplemented with complete protease inhibitor cocktail (Roche, 04693132001). DNA was sheared to an average fragment size of 250 bp using a Bioruptor (Diagenode) at 4 °C in polystyrene tubes, as confirmed by agarose gel electrophoresis. Cellular debris was removed by centrifugation. For each ChIP reaction, 2 mL of lysate was adjusted to a DNA concentration of 20 ng/μL (based on A₂₆₀ measurements), incubated with anti-FLAG M2 magnetic beads (Sigma, M8823) for 3 hours at 4 °C with end-over-end rotation. The beads were subsequently washed twice with 1 mL lysis buffer, once with lysis buffer containing 500 mM NaCl, and once with wash buffer (10 mM Tris/HCl, pH 8.0, 100 mM LiCl, 1 mM EDTA, 0.5% sodium deoxycholate, and 0.5% Nonidet P-40). Bound complexes were eluted using elution buffer containing 3× FLAG peptide (20 mM HEPES, pH 7.5, 150 mM NaCl, 1% Triton X-100, 0.4 mg/mL 3× FLAG peptide) and subjected to decrosslinking overnight at 65 °C in the presence of 10 μg RNase A and 40 μg Proteinase K. DNA was purified by phenol-chloroform-isoamyl alcohol extraction followed by precipitation with isopropanol. The pellet was washed twice with 75% ethanol, air-dried, and dissolved in 20 μL TE buffer. Library construction is performed by E-GENE Biotech Inc. (Shenzhen, China), and it was quantitatively and qualitatively detected using the Qubit® dsDNA HS Assay Kit and Agilent 2100 bioanalyzer. Finally, paired-end sequencing is conducted on the Illumina HiSeq PE150 platform (Illumina, San Diego, CA, USA). Quality control of raw fastq reads was conducted using FastQC v0.12.1 and trimming with Trimmomatic v0.38. Reads were aligned to the *E. coli* MG 1655 reference genome (GCF_000005845.2) and plasmid expressing PANGU using Bowtie2 v2.3.4.1.

### Fluorescence Microscopy

Single colonies of *E. coli* MG1655 harboring the pET28a plasmid with Pag1A-GFP and Pag1B-mcherry fusion expression or Pag2A-GFP fusion expression and Pag2B was cultured overnight in LB medium supplemented with 50 μg/mL kanamycin. Overnight cultures were diluted 1:100 into fresh LB medium containing 50 μg/mL kanamycin and 0.1 mM IPTG. Cultures were incubated at 37°C with shaking at 200 rpm. Uninfected control samples were collected at mid-exponential growth phase (OD600 = 0.5) using centrifugation (3,000 × g, 2 min). For phage infection assays, bacterial cultures were challenged with T4 using a MOI of 10, followed by 15-minute incubation at 37°C. Subsequently, cells were pelleted via centrifugation (3,000 × g, 2 min) and subjected to a single wash with ice-cold PBS. Cell pellets were treated with 0.02% Triton X-100 in PBS for 5 minutes to permeabilize membranes, followed by staining with 1 μg/mL DAPI (MCE, HY-D1738) in PBS for 15 minutes in the dark to label genomic DNA. After washing with PBS, cells were immobilized on poly-L-lysine-coated (0.1%) glass slides and imaged using a STELLARIS STED confocal microscope (Leica Microsystems) equipped with 405 nm (DAPI) and 488 nm (GFP) laser lines. Fluorescence images were analyzed using Leica Application Suite X (LAS X). Colocalization analysis between Pag1A/Pag2A (GFP channel) and genomic DNA (DAPI channel) was performed using ImageJ. Regions of interest were drawn across cells, and fluorescence intensity profiles were generated via the Plot Profile function. Overlapping peaks in the green (GFP) and blue (DAPI) channels indicated colocalization. Three independent biological replicates were performed to ensure statistical validity.

### Electrophoretic Mobility Shift Assay (EMSA)

To assess the DNA-binding activity of Pag1A, electrophoretic mobility shift assays (EMSAs) were performed. Double-stranded DNA fragments of 200, 400, and 1000 bp for use in EMSAs were amplified from the *E. coli* MG1655 genome using a 5’-FAM-labeled forward primer (GTGCCAAACACCACGTAGGA) in conjunction with the reverse primers R1 (ACTCGTCTGATGGGTGCAGC), R2 (CAATCGCCCCGCACAGCGAA), and R3 (TACCTGATGAATTCACTCTACAACG). Serial dilutions of Pag1A (4, 8, 16, 32, or 64 μM) were incubated with 10 nM of the respective FAM-labeled dsDNA in EMSA buffer (20 mM HEPES, pH 7.5, 150 mM NaCl, 5% glycerol) for 30 minutes at 37℃. The reaction mixtures were then loaded onto pre-run 6% native polyacrylamide gels and electrophoresed at 100 V for 50 minutes in 0.5× TBE buffer. Following electrophoresis, the free dsDNA and protein-DNA complexes were visualized using a Tanon-5200 Multi instrument (Tanon Science & Technology Co. Ltd., Shanghai, China).

### Bioinformatic Identification of Nucleoid-Associated Protein Homologs

All bacterial proteins annotated with the Nucleoid-associated protein NdpA domain (InterPro entry IPR007358; n=10,062) were retrieved as representatives of the 37-kDa nucleoid-associated protein (NAP) family. Multiple sequence alignment was performed using Clustal Omega (v1.2.4) with 10 refinement iterations to optimize positional homology^39^. Profile hidden Markov models (HMMs) were subsequently constructed from the consensus alignment using hmmbuild (HMMER v3.4) with default parameters^40^.

The complete proteomes of 2,387,175 bacterial and 25,153 archaeal genomes (NCBI GenBank, accessed 2024-12-04) were systematically interrogated. Open reading frames (ORF) were predicted *de novo* using Prodigal (v2.6.3) in metagenomic mode (-p meta) to ensure sensitivity for atypical genomes^41^. HMMER3’s hmmsearch was implemented with domain-specific bit score thresholds corresponding to E-value < 1e-10, followed by length filtering (≥300 aa) based on empirical observations of characterized NAPs (mean length 333 aa).

### NAP operon Architecture and Defense System Association

Prokaryotic operon structures were delineated using a bidirectional scanning algorithm with a maximum intergenic distance threshold of 40-bp between ORF. Genomic regions flanking identified NAP genes were systematically interrogated until encountering intergenic regions exceeding this threshold, with all contiguous ORF annotated as co-transcribed units.

To mitigate assembly fragmentation artifacts, operons spanning contig termini (within 10 kb of contig edges) were excluded from downstream analyses. This conservative filtering retained only operons with ≥ 10 kb internal sequence context for reliable functional annotation.

Putative defense associations were identified using DefenceFinder (v1.2.2) with extended 10-kb flanking regions. To eliminate redundancy^42^, the DNA sequences of operons linked to defense systems were deduplicated using SeqKit (v2.1.0)^43^. Non-NAP proteins within defense-associated operons underwent hierarchical clustering via MMseqs2 v13.45111 (sensitivity 0.9, coverage 0.9)^44^.

Frequently occurring clusters (n ≥ 10 members) were retained for PANGU system classification. The function of non-NAP proteins in NAP operons were annotated using emapper (v2.1.12)^45^.

### Identification of PANGU systems

A detailed multiple sequence alignment (MSA) of homologous proteins from canonical PANGU systems was performed using MAFFT (v7.505) with 1000 iterations^46^. The resulting alignment was used to construct HMM models, which were subsequently employed for screening the NAP operons using hmmsearch (v3.4)^40^. This screening yielded a large number of results, with many proteins being hit multiple times due to shared similarities across different PANGU system types. To identify true, non-redundant PANGU systems, we merged all results, sorted them by gene position, and selected the hit with the smallest E-value for proteins appearing multiple times. Incomplete PANGU systems were discarded, and 10 high-quality PANGU system types were successfully identified within the operons.

### Phylogenetic Reconstruction and Taxonomic Analysis

To mitigate branch length artifacts arising from concatenated multi-protein analyses of diverse PANGU systems, we restricted phylogenetic reconstruction to conserved NAP sequences as system-type proxies.

Standalone NAPs without operonic defense associations underwent flanking region analysis (10-kb upstream/downstream screening) followed by MMseqs2 clustering (70% identity threshold) to select representative sequences.

Phylogenetic trees were constructed using both rapid topology approximation and model-optimized reconstruction methods. The rapid topology approximation employed MUSCLE (v5.3) with the “--super5” parameters for MSA and FastTree (v2.1.11) for tree construction^47,48^, providing a preliminary assessment of evolutionary relationships. The model-optimized reconstruction involved more precise alignment using MAFFT with the --localpair option, followed by phylogenetic tree construction with IQ-TREE (v2.4.0) using the best-fit model Q.yeast+F+R8 and 1500 bootstrap iterations to improve tree accuracy^49^. The resulting trees were visualized using iTOL (v6)^50^.

Genomes harboring NAP homologs were taxonomically profiled via GTDB-Tk (v2.4.0)^51^. This enabled automatic species classification and facilitated the assessment of species distribution across phyla.

### Co-evolutionary Analysis of Pag1A and Pag1B

To explore the evolutionary relationship between Pag1A and Pag1B, both sequence-based phylogenetic correlation and structural interaction prediction were performed. Evolutionary distances between Pag1A/Pag1B homolog pairs and their corresponding host species were calculated with Pearson correlation coefficients (r) computed to assess phylogenetic congruence. Phylogenetic trees for both proteins were reconstructed using the method described previously: MAFFT v7.505 followed by IQ-TREE v2.4.0 under the Q.yeast+I+R8 (Pag1A) and Q.pfam+F+I+R7 (Pag1B) model with 1500 ultrafast bootstrap replicates.

For host species phylogeny reconstruction, genomes containing Pag1 homologs were analyzed using GTDB-Tk v2.4.0 with the Bac120 marker set. A concatenated alignment of 120 bacterial marker genes was subjected to partitioned phylogenetic analysis in IQ-TREE (Q.pfam+R9 model selected by ModelFinder, 1500 ultrafast bootstraps).

Structural co-evolution was assessed by predicting 3D conformations for 10 phylogenetically diverse Pag1A/Pag1B pairs using AlphaFold3. Pairwise comparisons of these predicted structures were made to predict potential protein-protein interactions, and the reliability of these predictions was assessed using the ipTM score. This analysis provided insights into the structural co-evolution of Pag1A and Pag1B proteins.

### Statistics Analysis

For all statistics analysis, the assays were performed in three independent replicates as indicated in the figure legends, and the unpaired t test was used to calculate P values.

## Data and materials availability

Data are publicly available. The atomic coordinates for the cryo-EM structure of the Pag1A-Pag1B complex (PDB 9UFP) were deposited to the PDB (www.rcsb.org). The cryo-EM density maps reported in this study for the Pag1A-Pag1B complex (EMD- 64120) was deposited to the EM Data Bank. All additional data reported in this paper will be shared by the lead contact upon request. This paper does not report original code.

## Acknowledgments

We thank Dr. Xiaolan Zhang and Dr. Yao Wu at the Institute of Microbiology for providing technical support.

## Author contributions

M.L. conceived and supervised the project with valuable suggestions from B.L., S.Z., X.S., and V.H.. M.L., B.L., X.S., and Z.L. designed the experiments. Z.L. performed bacteriophage-host interaction assays, protein-protein interaction assays and enzyme activity characterization. F.C., J.D., I.T., H.Y. and J.G. performed quantitative analysis of phage plaque efficiency. B.L., S.Y., Z.Y. and J.W. performed protein physicochemical property identification and structural elucidation. X.S., G.A. and A.G.P. performed LC-MS analysis. S.Z. and J.M. performed bioinformatics-driven computational analysis. M.L. and Z.L. drafted the manuscript. M.L., V.H., B.L., and S.Z. finalized the manuscript with input from all authors.

## Extended Data

**Extended Data Fig. 1.**
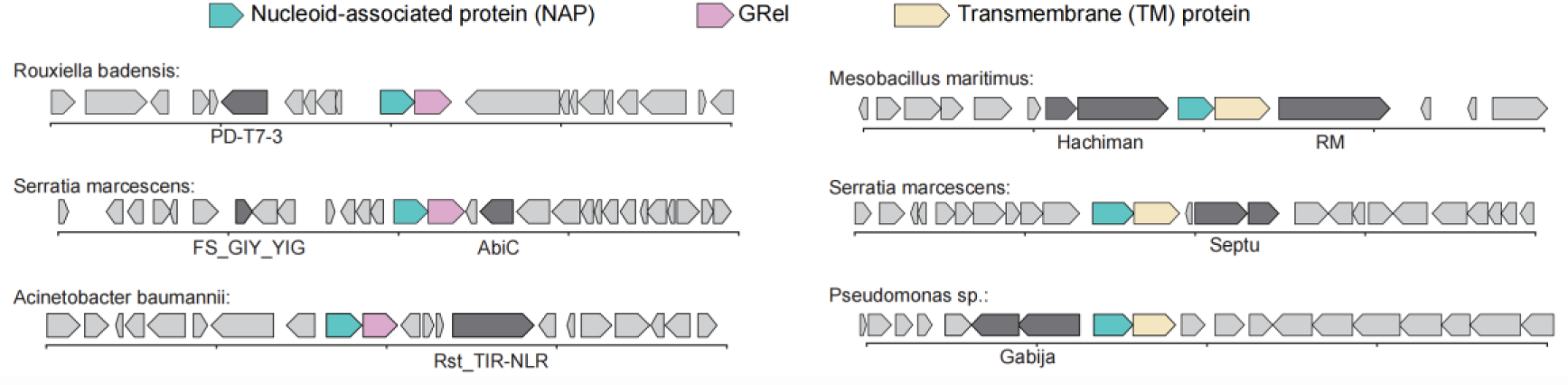
Two conserved NAP-encoding operons (putative PANGU systems) co-localized with known defensive genes (dark grey). For each type of NAP-encoding operon, three representative examples and their genomic contexts are shown. RM, restriction-modification.

**Extended Data Fig. 2.**
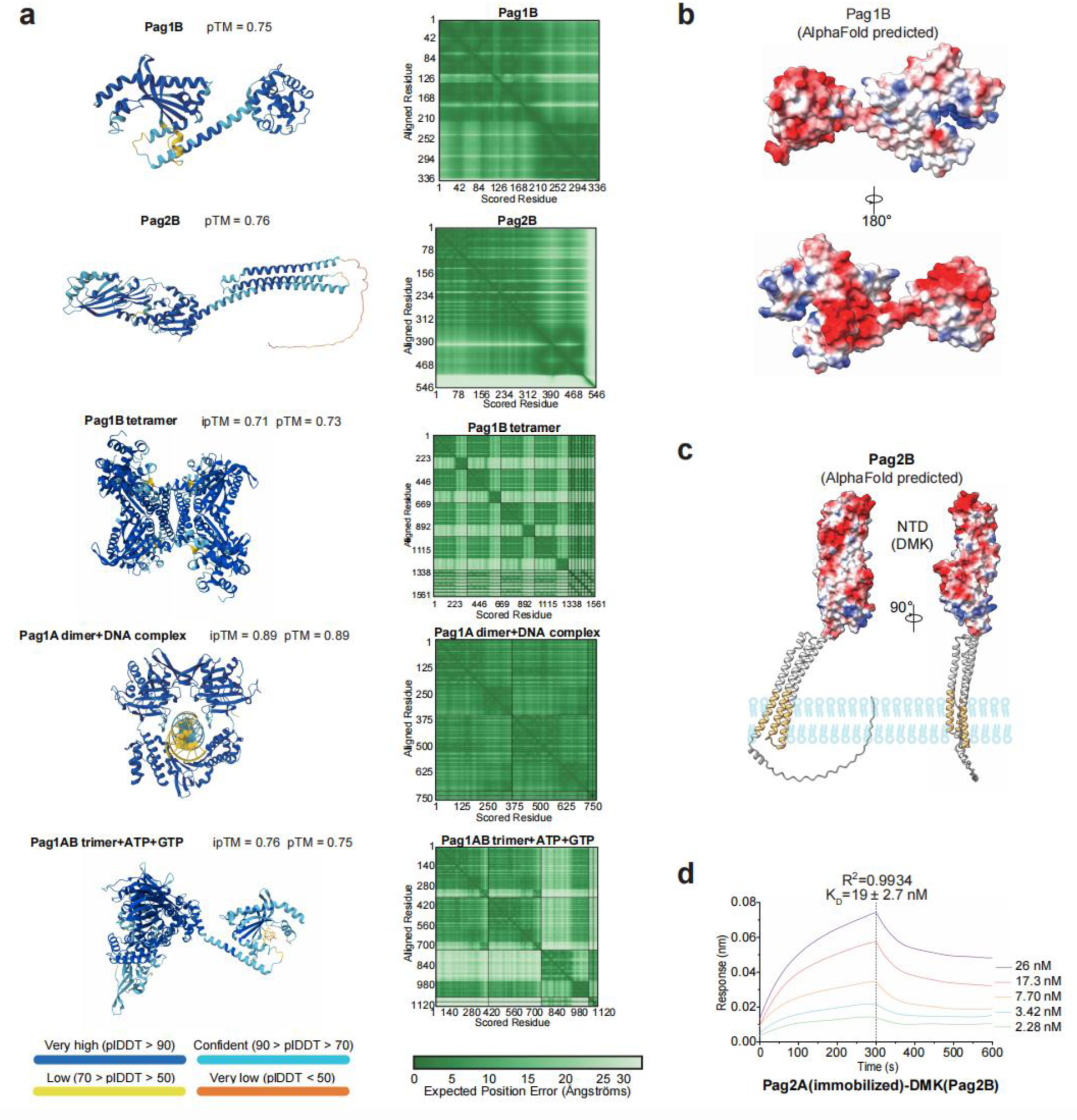
Structural predictions of Pag1B, Pag2B, and Pag1A. (a) Per-residue pLDDT confidence scores mapped onto the predicted structures for Pag1B, Pag2B, Pag1B tetramer, and DNA-bound Pag1A, with the predicted aligned error (PAE) between residues given (right). (b) The surface charge distribution of Pag1B was visualized by mapping electrostatic potentials, with surface atoms colored on a red-to-blue scale representing negative to positive potentials, respectively. (c) An electrostatic potential map of Pag2B is given to show its DMK domain. Light yellow denotes transmembrane region at the C-terminus. The structures are displayed with a 90 ° rotational offset along the z-axis to highlight structure and electrostatic distribution. (d) Bio-Layer Interferometry (BLI) analysis of Pag2A and the DMK domain of Pag2B. Serial dilutions of the DMK domain of Pag2B were injected onto Pag2A-loaded biosensors for a 300-second association phase, followed by a 300-second dissociation phase.

**Extended Data Fig. 3.**
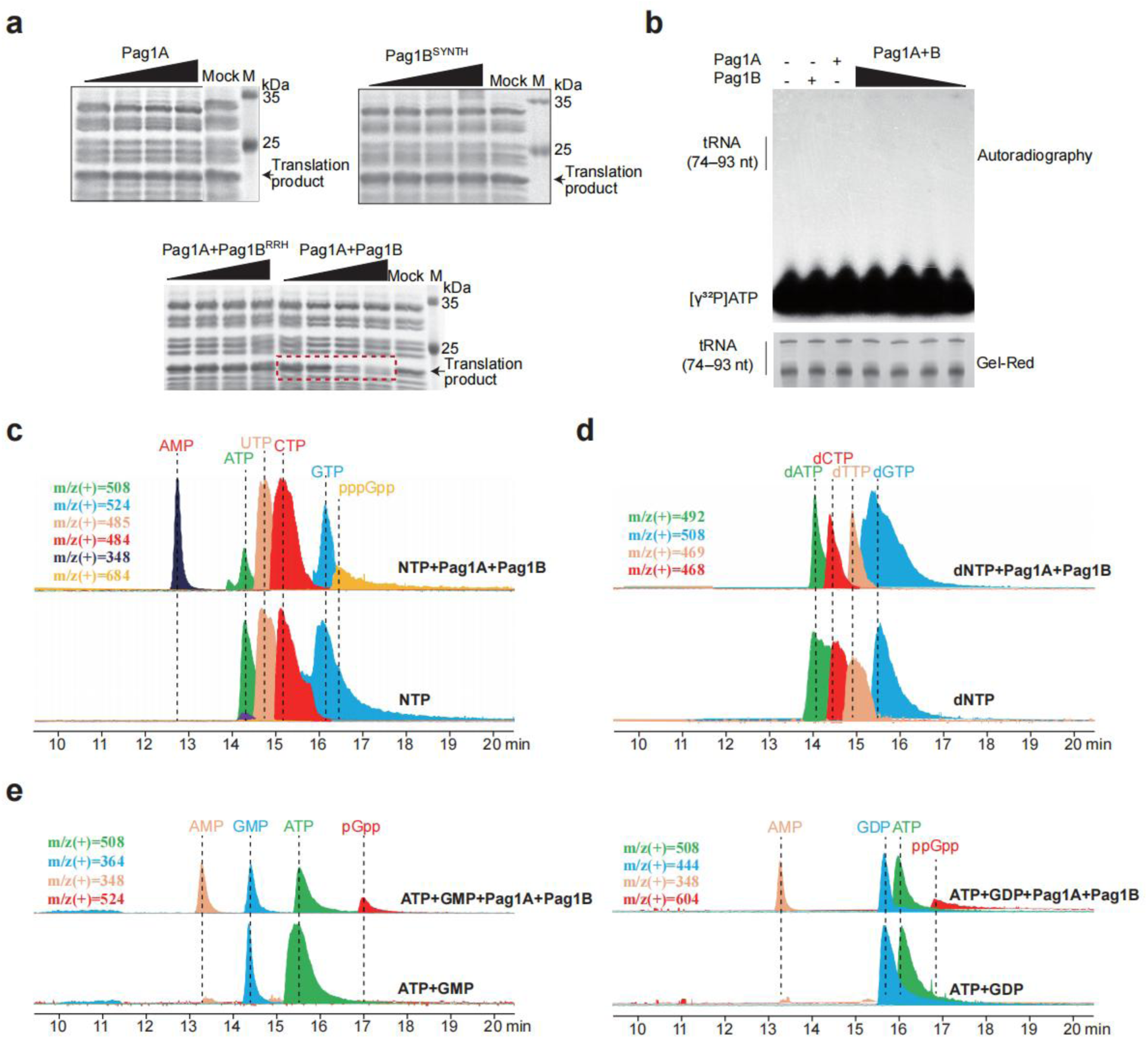
*In vitro* translation inhibition and enzymatic activity assays. (a) Translation inhibition by Pag1A alone, the isolated SYNTH domain of Pag1B, or Pag1A plus the Pag1B^RRH^ triple mutant. *In vitro* transcription-translation assays monitored DHFR production from its DNA template. Purified protein was assayed at concentrations of 62.5, 125, 250, or 500 nM, individually or in combination. (b) Autoradiography of *E. coli* bulk tRNA after incubation with Pag1A and/or Pag1B in the presence of [γ^32^P] ATP. GelRed staining of the same gel confirms equal tRNA loading. Representative images from three biological replicates are shown. (c**–**e) LC-MS analysis of nucleotide substrates after incubation with purified Pag1A and Pag1B. (c) NTP mixture; (d) dNTP mixture; (e) GMP and GDP. Extracted-ion chromatograms are shown with m/z values indicated.

**Extended Data Fig. 4.**
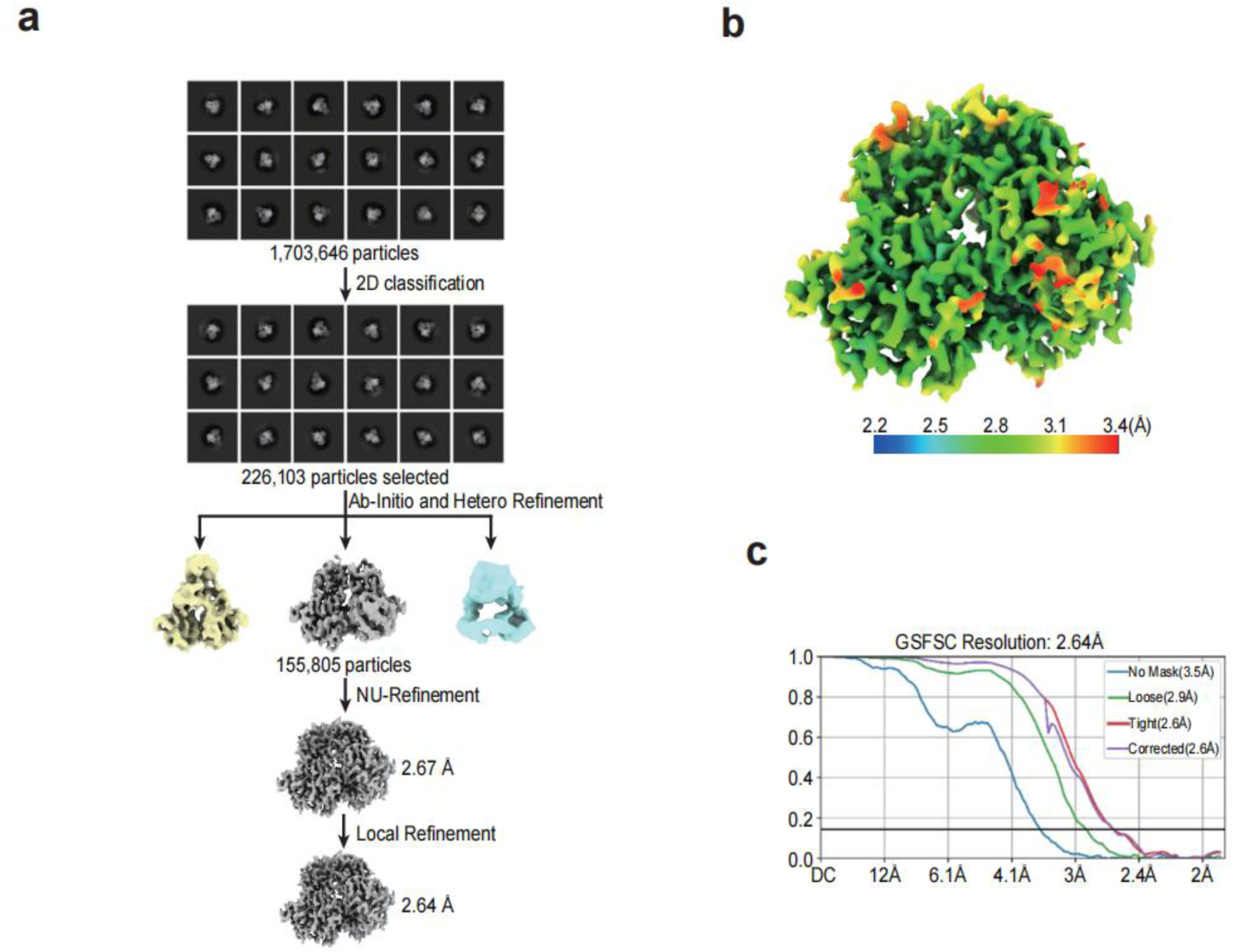
Structure determination of the Pag1A-Pag1B heterotrimer. (a) Cryo-EM processing pipeline. Schematic representation of the image processing pipeline used to solve the structure of the Pag1A-Pag1B heterotrimer complex. (b) Local resolution estimation mapped onto the final cryo-EM density map of the Pag1A-Pag1B complex, highlighting regional variations in resolution quality. (c) Fourier shell correlation (FSC) curve derived from two independently refined half-maps. The gold-standard resolution cutoff (FSC = 0.143) is indicated with a solid line.

**Extended Data Fig. 5.**
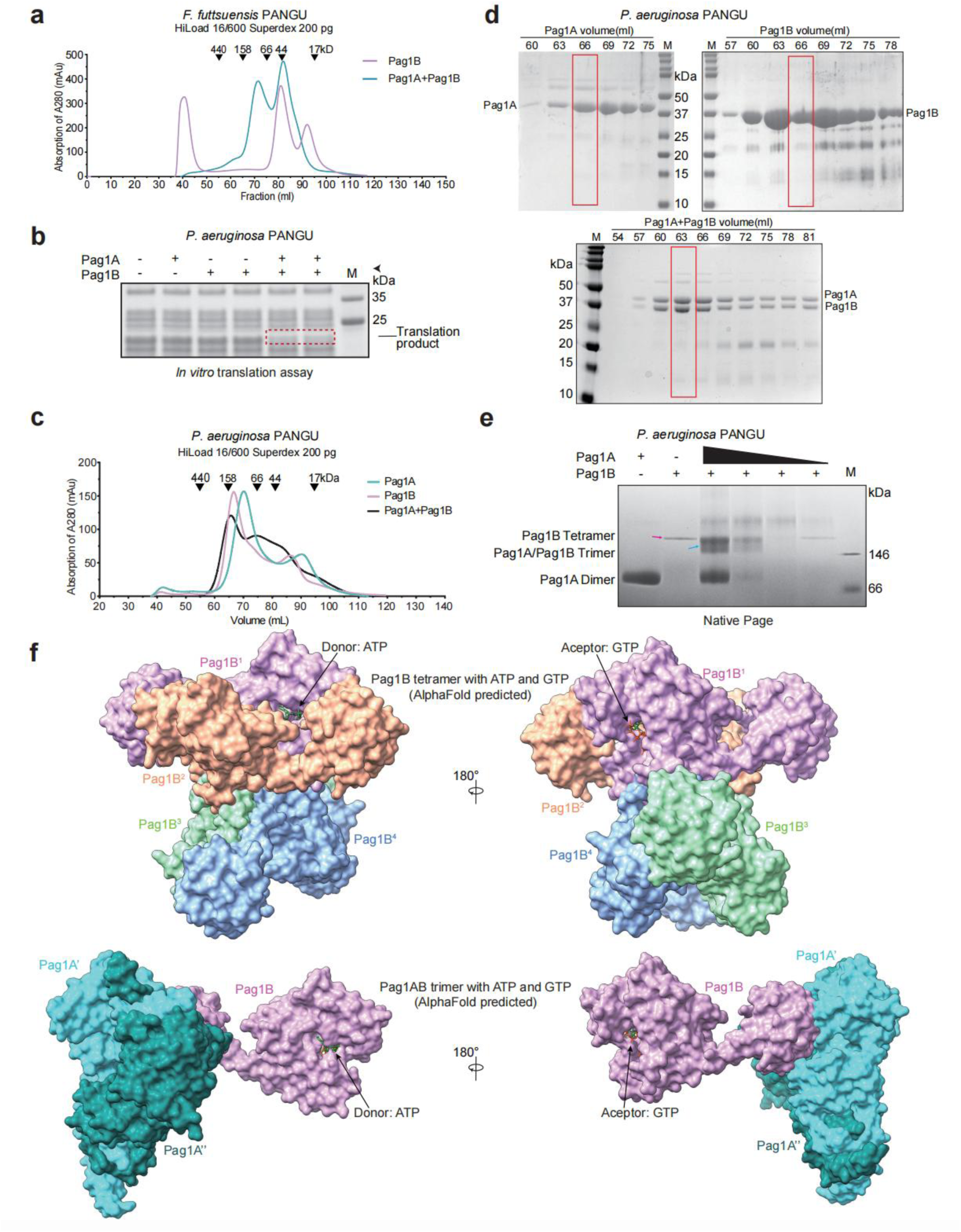
Pag1B forms a tetramer that can be disassembled by Pag1A. (a) SEC profiles of *F. futtsuensis* Pag1B in the presence or absence of cognate Pag1A. The elution positions of standard protein markers are marked above the trace. (b) Translation inhibition by the *P. aeruginosa* PANGU system. *In vitro* transcription-translation assays monitored DHFR production from its DNA template. Purified Pag1A and Pag1B was assayed at concentrations of 500 nM, separately or in combination. (c) SEC analysis of *P. aeruginosa* Pag1A, Pag1B, or their 1:1 mixture; standards as in (a). (d) SDS-PAGE of selected SEC fractions. Numbers indicate elution positions. Fractions subjected to SEC-MALS are boxed in red. The Pag1B band (red box) was intentionally under-loaded (one-third of adjacent samples). M, protein markers in kilodaltons (kDa). (e) Native PAGE analyzing showing Pag1A-induced dissociation of the Pag1B tetramer. (f) Surface comparison of the AlphaFold-modeled Pag1B tetramer:ATP:GTP complex versus the Pag1AB trimer:ATP:GTP complex (see Extended Data Fig. 2a). Pag1B tetramerizes as a head-to-tail dimer of dimers. Donor ATP and acceptor GTP are represented as green and orange sticks, respectively.

**Extended Data Fig. 6.**
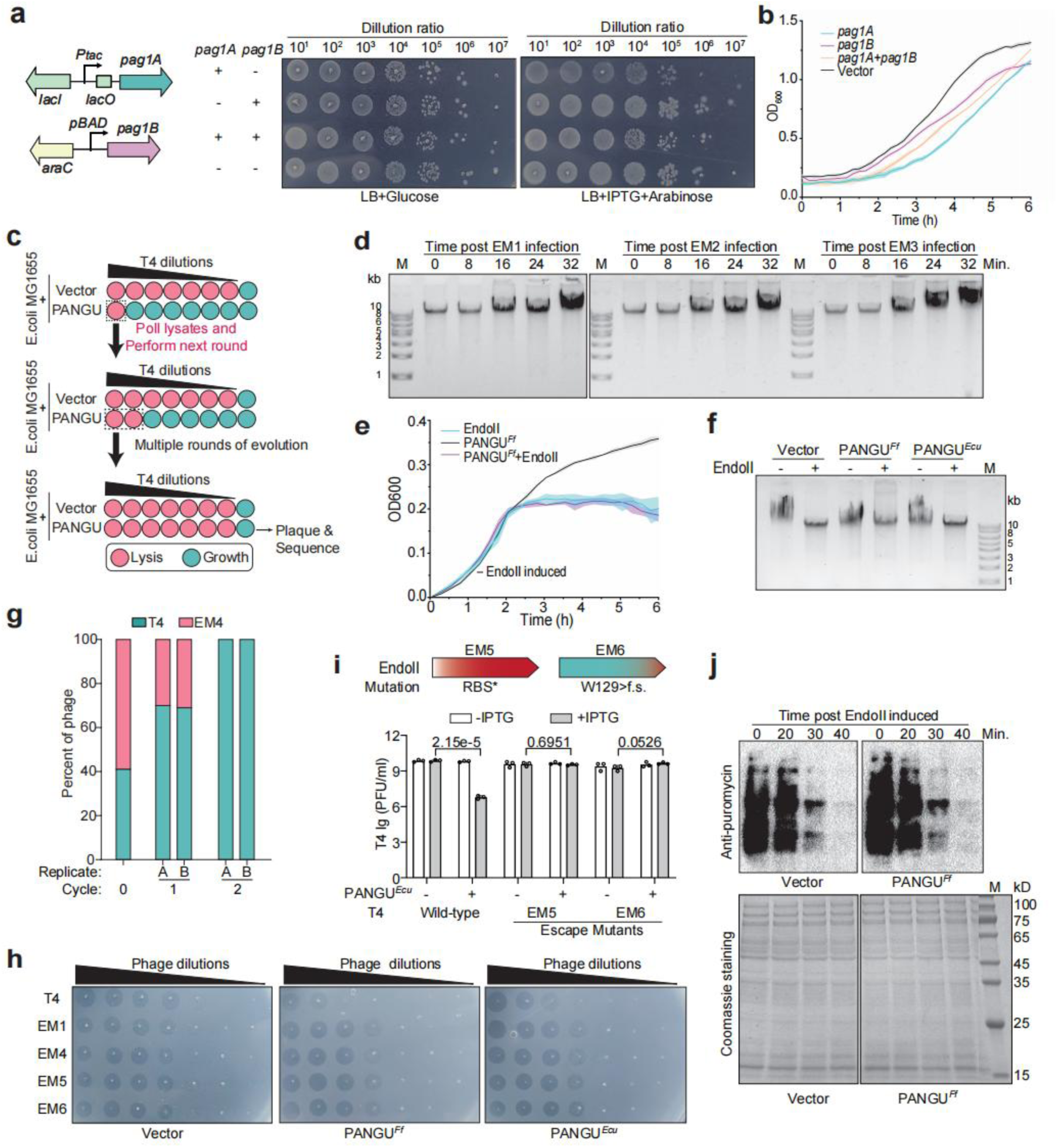
Screening for and analysis of T4 phage-triggered PANGU defense activation. (a) Schematic of inducible expression setup for Pag1A and Pag1B. Serial dilution and plating of *E. coli* cells on inducing or non-inducing plates, using empty vectors (-) as controls. Representative images from three biological replicates are shown. (b) Growth curves of *E. coli* cells expressing Pag1A and Pag1B individually or in combination. Cells with empty vectors served as controls. Data points indicate mean ± SD from three biological replicates. (c) Schematic of the experimental evolution approach to screen for T4 mutants capable of evading PANGU defense. Pink wells indicate phage-induced cell lysis, while cyan wells indicate bacterial growth. (d) Agarose gel electrophoresis of genomic DNA extracted from *E. coli* cells infected by EM1, EM2, or EM3. (e) Growth curves of *E. coli* cells expressing EndoII and/or PANGU*^Ff^*. IPTG was used for induction. Data points represent the mean ± SD from three biological replicates. (f) Agarose electrophoresis analysis of genomic DNA extracted from PANGU-expressing *E. coli* cells with or without *denA* expression. (g) Competition assay between T4 and EM4 on *E. coli* cells. Phages were mixed at a 1:1 ratio and used to infect bacteria at an MOI of 0.01. Two cycles of competition were performed, and phage mixtures were sequenced to quantify T4 and EM4 fractions before and after each cycle. The *y*-axis shows the relative abundance of each phage, and the *x*-axis indicates the competition cycle. Initial mixtures before competition are labeled as cycle 0. Data from two independent replicates (A and B) are shown. (h) Serial tenfold dilutions of T4, EM1, EM4, EM5, or EM6 phages spotted on lawns of *E. coli* cells expressing PANGU*^Ff^* or PANGU*^Ecu^*. EV, empty vector. Representative images from three biological replicates are shown. (i) PFUs of T4, EM5, or EM6 infecting *E. coli* cells expressing the PANGU*^Ecu^*. Cells with the empty vector served as a control (-). IPTG was used for induction. Data represent the mean ± standard deviation (SD) of three biological replicates. Mutations in the *denA* gene of EM5 and EM6 are shown above. f.s. : frame shift. (j) Translation efficiency of *E. coli* cells expressing both PANGU*^Ff^* and *denA*. Cells were collected at different time points post *denA* induction. Translation efficiency was assessed by puromycin incorporation and immunoblotting with anti-puromycin antibodies. Coomassie staining of the same gel indicates equal protein loading. Representative images from two independent biological replicates are shown.

**Extended Data Fig. 7.**
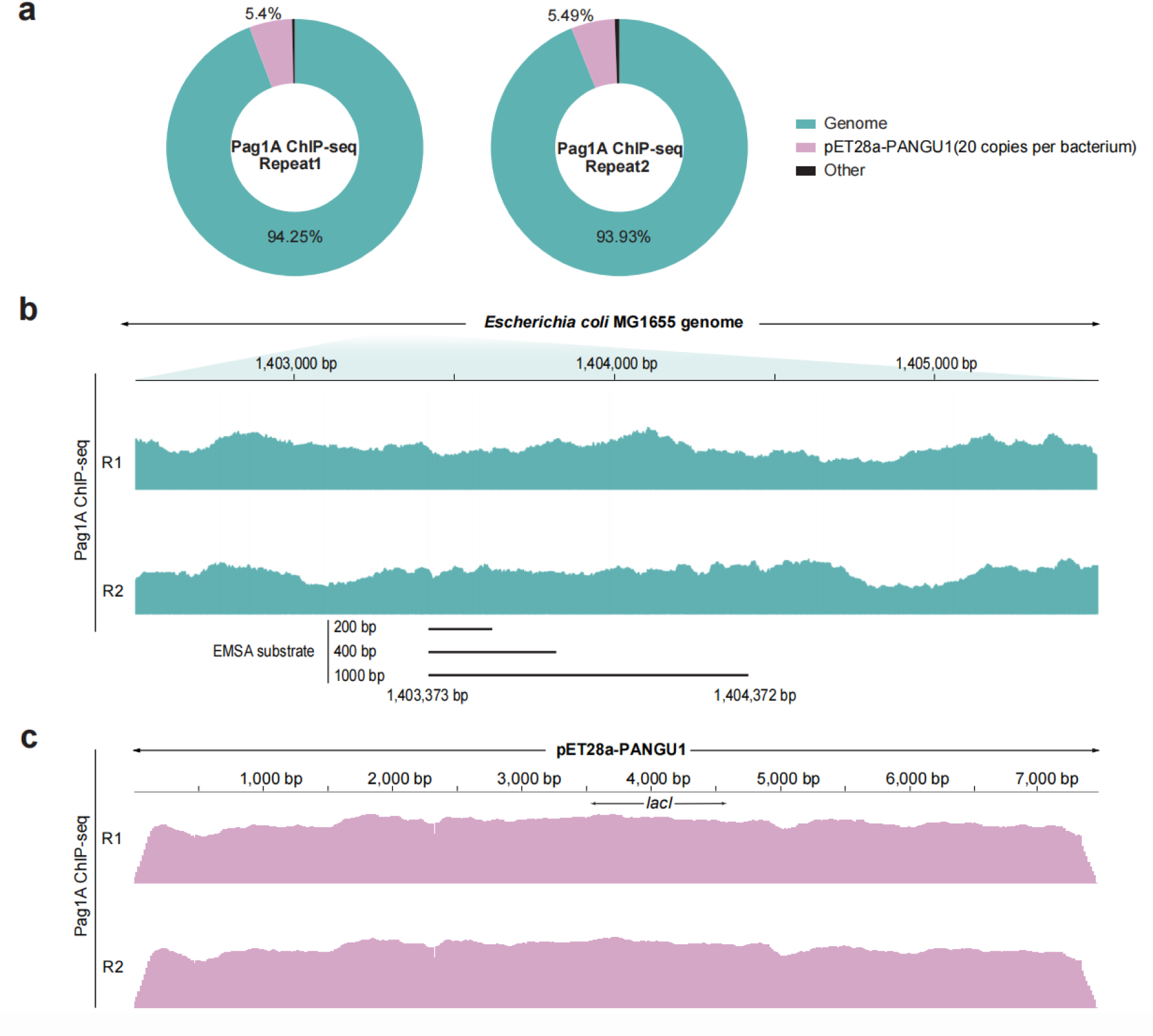
ChIP-seq analysis of Pag1A. (a) Percentage of Pag1A ChIP-seq reads mapping to the *E. coli* MG1655 chromosome versus the PANGU *^Ff^* -expressing pET28a plasmid. (b) Read density across a representative 3-kb chromosomal window. The fragments used for the EMSA assays (Fig. 5b, d) are indicated below the trace. (c) Read coverage along the PANGU*^Ff^*-pET28a plasmid.

**Extended Data Fig. 8.**
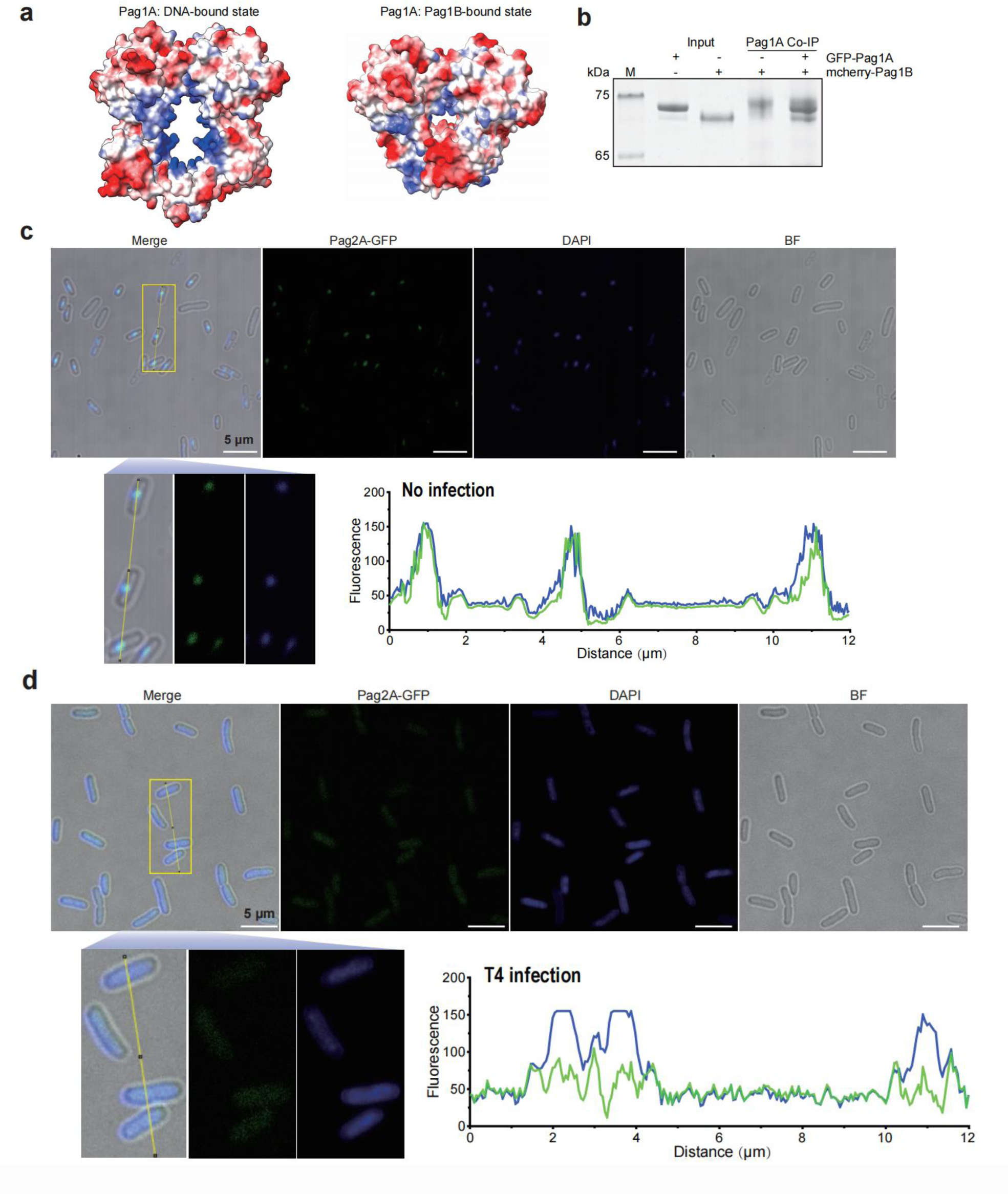
PagA-DNA colocalization analysis. (a) Electrostatic surface potentials of the Pag1A dimer in the DNA-bound and the Pag1B-bound states. (b) Pull-down of Pag1B (N-terminal mcherry-tagged) by Pag1A (N-terminal GFP-tagged). M, protein markers in kilodaltons (kDa). (c, d) Fluorescence microscopy of PANGU*^Ecu^*-expressing *E. coli* cells under uninfected and T4-infected conditions. Genomic DNA (DAPI stained) and Pag2A (GFP-fused) were visualized using the blue and green channels, respectively. Merged images were analyzed using ImageJ’s Plot Profile tool to quantify signal correlation, shown on the right. Scale bar, 5 μm.

**Extended Data Fig. 9.**
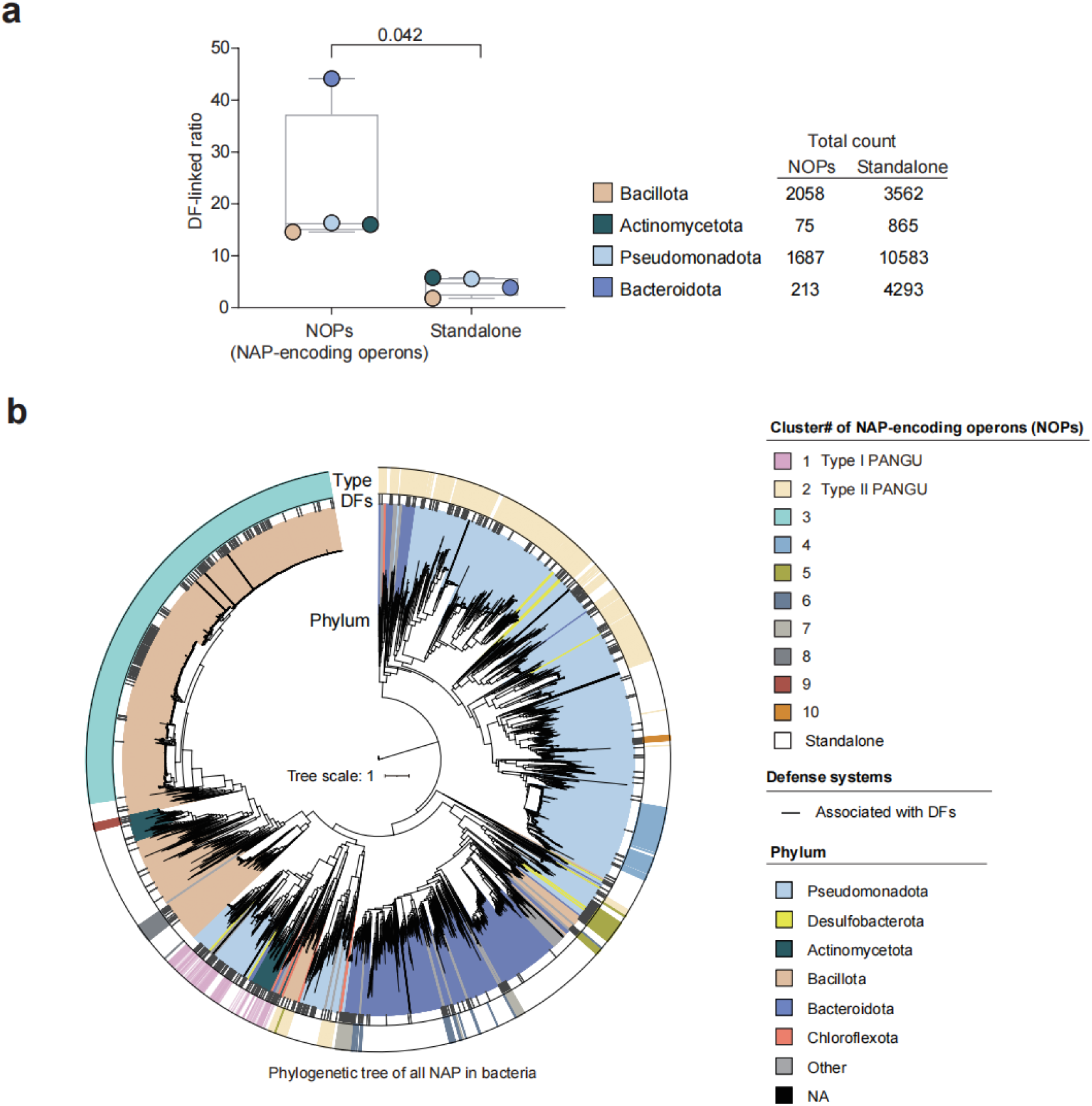
Phylogenetic analysis of standalone and operon-encoded NAPs. (a) Box plot showing the proportion of standalone NAPs or NAP-encoding operons linked to known defense (DF) genes across different bacterial phyla. Total numbers of standalone NAPs and NOPs per phylum are listed on the right. The *P* value was calculated by a two-sided *t*-test. (b) Phylogenetic tree of standalone NAPs and NAPs from the 10 NAP-encoding operon (NOP) clusters. From the inner to outer circles: (1) phylum of NAP-containing strains (color-coded); (2) genetic linkage to known defense genes (DFs); (3) representation of standalone NAPs (white) or operon-encoded NAPs (color-coded).

**Extended Data Fig. 10.**
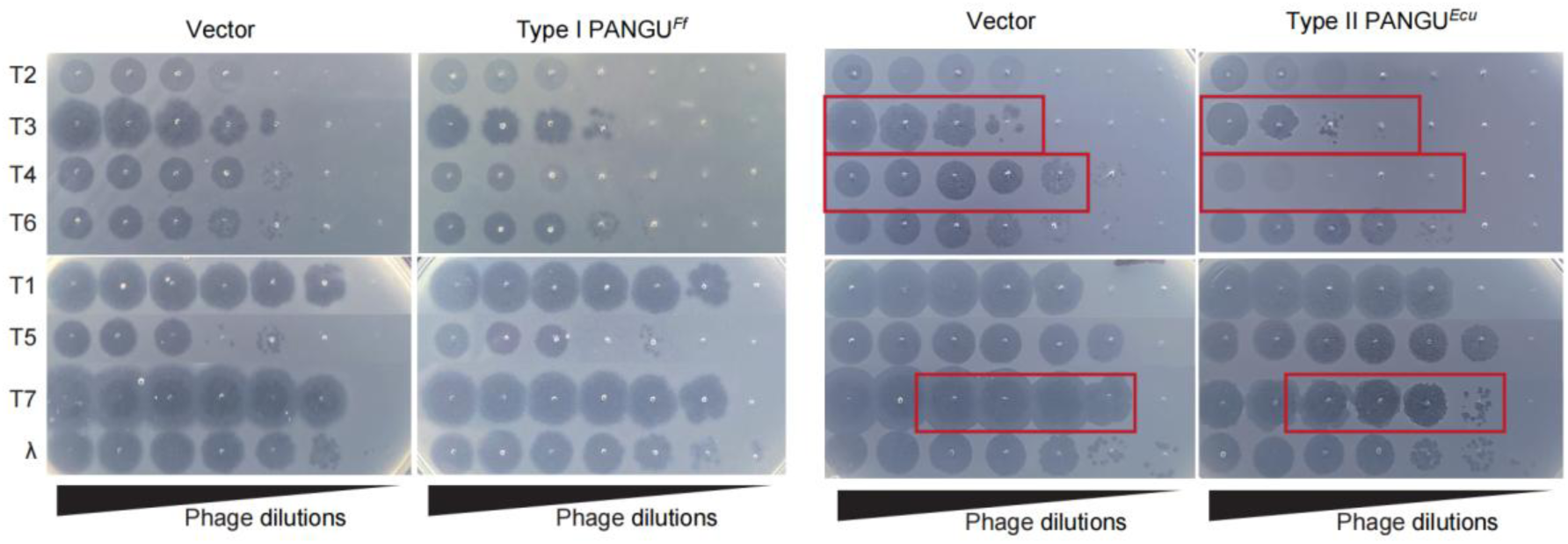
Anti-phage activity of PANGU*^Ff^*/PANGU*^Ecu^* against different coliphages. Serial, tenfold dilutions of coliphages spotted on lawns of *E. coli* cells expressing PANGU*^Ff^*or PANGU*^Ecu^*. Cells harboring the empty vector served as a control. Representative images from three biological replicates are shown.

**Extended Data Fig. 11.**
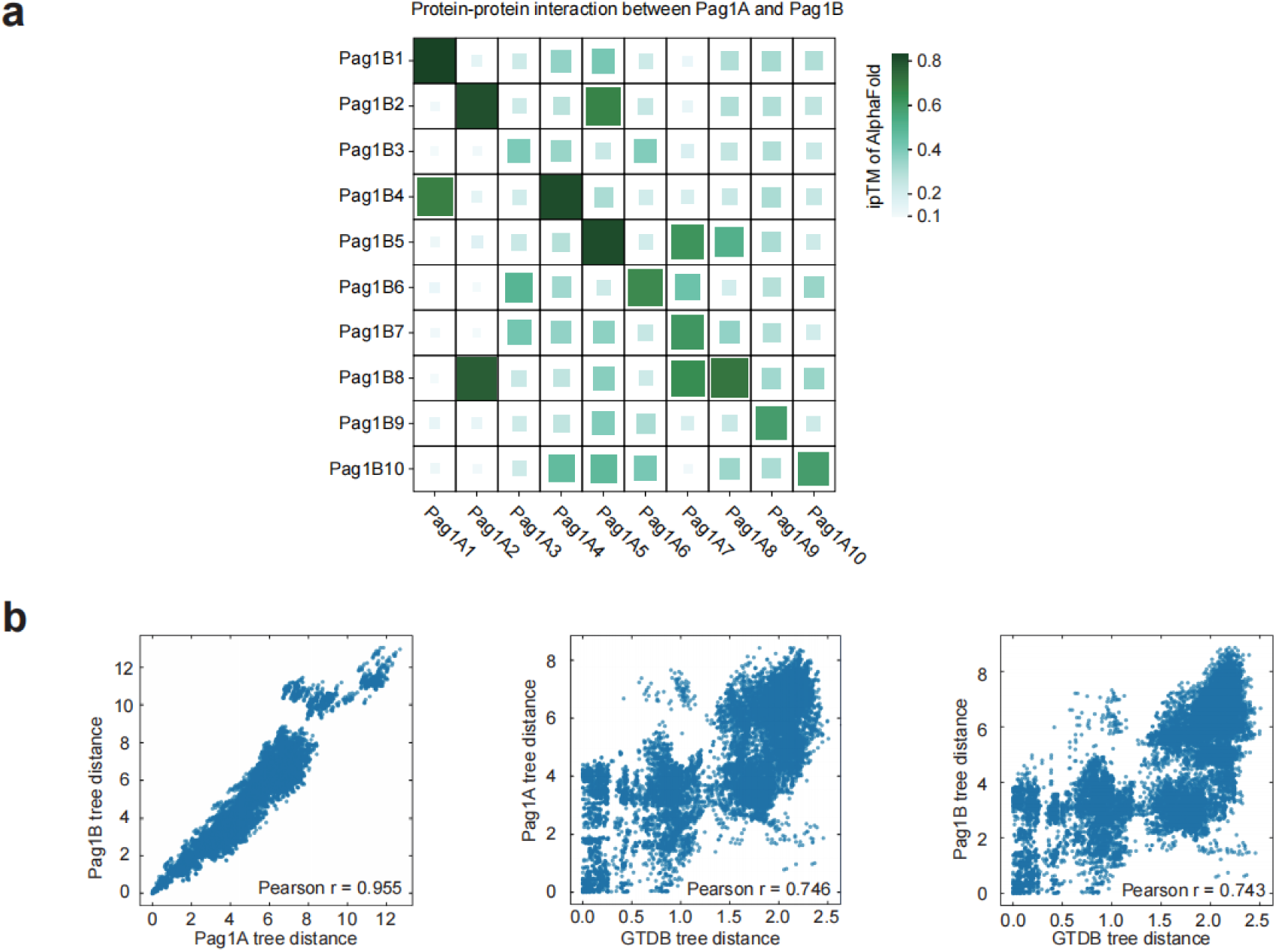
Interaction specificity and co-evolution analysis of Pag1A and Pag1B. (a) AlphaFold3-generated heatmap showing the interface predicted template modeling (ipTM) scores for the interactions between Pag1A and Pag1B proteins from different species. (b) Pearson correlation analysis comparing evolutionary distances between Pag1A, Pag1B, and their host species phylogenetic trees. Each point represents the pairwise patristic distance between two shared leaf nodes computed from the two phylogenetic trees. The x-axis and y-axis represent the evolutionary distances of protein or species trees, respectively.

**Extended Data Table 1.**
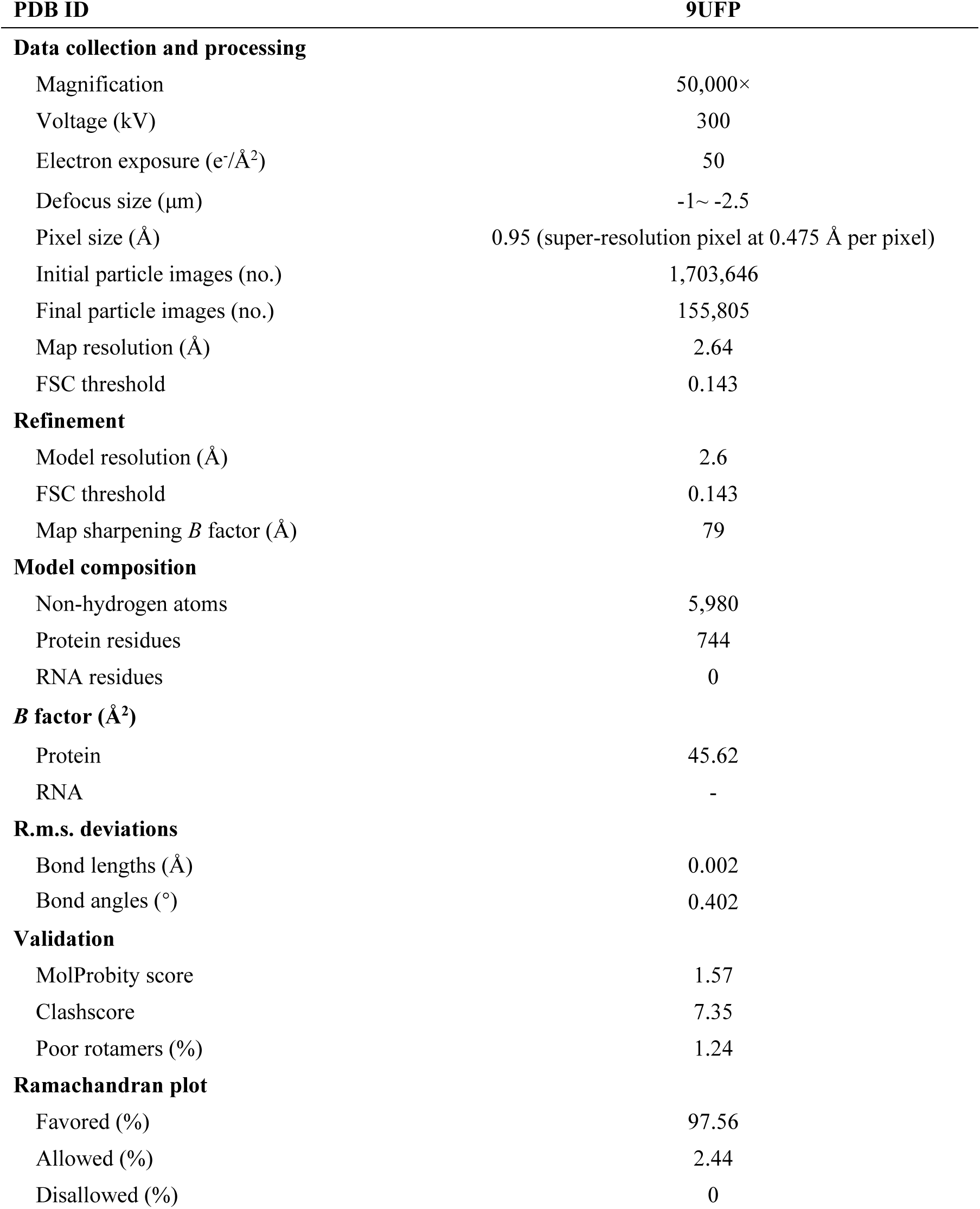
Cryo-EM data collection, refinement and validation statistics.

